# RNA-CLAMP Enables Photo-activated Control of CRISPR-Cas9 Gene Editing by Site-specific Intramolecular Cross-linking of the sgRNA

**DOI:** 10.1101/2021.04.22.441030

**Authors:** Dongyang Zhang, Shuaijiang Jin, Luping Liu, Ember Tota, Zijie Li, Xijun Piao, Neal K. Devaraj

**Author notes:** **Corresponding Author**, Neal K. Devaraj.

## Abstract

Here we introduce RNA-CLAMP, a technology which enables site-specific and enzymatic cross-linking (clamping) of two selected stem loops within an RNA of interest. Intramolecular clamping of the RNA can disrupt normal RNA function, whereas subsequent photo-cleavage of the crosslinker restores activity. We applied the RNA-CLAMP technique to the single guide RNA of the CRISPR-Cas9 gene editing system. By clamping two stem loops of the single-guide RNA (sgRNA) with a photo-cleavable cross-linker, gene editing was completely silenced. Visible light irradiation cleaved the crosslinker and restored gene editing with high spatiotemporal resolution. Furthermore, by designing two photo-cleavable linkers which are responsive to different wavelength of lights, we achieved multiplexed photo-activation of gene editing in mammalian cells. Notably, although the Cas9-sgRNA RNP is not capable of DNA cleavage activity upon clamping, it maintained the capability to bind to the target DNA. The RNA-CLAMP enabled photo-activated CRISPR-Cas9 gene editing platform offers clean background, free choice of activation wavelength and multiplexing capability. We believe that this technology to precisely and rapidly control gene editing will serve as a versatile tool in the future development of stimuli responsive gene editing technologies. Beyond gene editing, RNA-CLAMP provides a site-specific tool for manipulating the internal structure of functional RNAs.

## Introduction

Many creative approaches have been developed to study and control the function of RNAs. RNA probes most often rely on non-covalent interactions, such as in the use of anti-sense DNA oligos, MS2-tagging, or RNA aptamers [1a, 1b]. Compared to techniques relying on non-covalent interactions, covalent RNA modification strategies offer an additional level of robustness, which can be especially important for less abundant RNA in harsh cellular conditions. However, due to the complex structure of RNA, as well as its relative instability, few approaches have been developed which allow for site-specific and covalent modification of RNA [1c–1e]. Chemical cross-linking is commonly used for studying RNA-protein and RNA-nucleic acid interactions [2a–2b]. Crosslinking of RNA enables rapid identification of potential regions of inter- and intramolecular interactions as well as study of the three-dimensional structure of complex RNA [2c–2e]. However, to date, no method has been established to achieve site-specific and covalent cross-linking of two user-defined sites within an RNA of interest. Many RNAs, such as the single-guide RNA (sgRNA) of the CRISPR-Cas9 gene editing system, require secondary and tertiary structure for function. Therefore, the ability to site-specifically cross-link two internal nucleotides within an RNA of interest could become an extremely valuable technology to study and manipulate RNA function. Here, we report an RNA-modifying technique, termed RNA-CLAMP, which allows for site-specific and post-transcriptional cross-linking of two specific guanine nucleotides within a single RNA of interest. We applied the RNA-CLAMP technique to the sgRNA of CRISPR-Cas9 and achieved optical control of gene editing in mammalian cells with high spatiotemporal resolution and multiplexing capability.

The RNA-guided CRISPR-Cas9 gene editing system has revolutionized biomedical research and numerous translational applications are being explored [3a–3f]. Conditionally activated CRISPR-Cas9 gene editing systems allow for greater gene editing precision by limiting Cas9-mediated DNA cleavage to a specific time and location. Such techniques offer reduced levels of off-target genome modification as well as greater spatiotemporal resolution to study complex gene networks and DNA repair processes [4a–4b]. Multiple methods have been established to chemically control CRISPR-Cas9 gene editing. For example, small-molecule inducers, such as doxycycline and rapamycin were used to induce gene editing in mammalian cells [5a–5c]. Conditional activation of CRISPR-Cas9 gene editing can also be achieved via transient delivery of purified Cas9-gRNA complex [6–8]. However, these chemical approaches have limitations. First, small-molecule inducers, for example rapamycin, can cause undesirable biological effects and toxicity. Second, the slow diffusion rate of small-molecule inducers makes the regulation of gene editing less responsive and precise. On the other hand, optical control of CRISPR-Cas9 offers non-invasive manipulation of gene editing with excellent spatiotemporal resolution. By controlling the irradiation time period and light intensity, cellular toxicity can be minimized. Moreover, different wavelength of lights might be used to trigger gene editing at multiple genomic loci, enabling multiplexed activation of gene editing.

Several approaches have been recently developed to achieve optical control of CRISPR-Cas9 gene editing by focusing on modifying either the Cas9 protein or the gRNA. Protein centric approaches aim to modify key amino acid residues within Cas9 to control protein function. For example, the Deiters lab has developed a genetically encoded light-activated Cas9 by engineering caged lysine amino acids within the Cas9 protein [9]. Alternative approaches include the use of split Cas9 proteins fused to a pair of photo-dimerizing proteins, to achieve optical control of CRISPR-Cas9 gene editing [10]. Some of these approaches suffer from incomplete caging and compromised activity when uncaged. Instead of modifying the Cas9 protein, RNA centric approaches focus on modifying the gRNA to optically control CRISPR-Cas9 gene editing. A typical strategy uses photo-cleavable DNA oligonucleotides that are complementary to the 20-nt target region of the gRNA to block its binding towards the target DNA [11]. The caged gRNA can be released upon photo-cleavage of the protector DNA oligo, resulting in activation of gene editing [11]. However, restricted by the chemical linker which was used to synthesize the protector DNA oligo, toxic UV light had to be applied to cells during activation. A more recent study applied engineered caged gRNA by substituting four nucleobases that are evenly distributed throughout the 5’-protospacer region with caged nucleobases during solid-state RNA synthesis [12]. This approach offers excellent activation rate and clean background at the caged stage. However, it requires solid-state synthesis, which is not widely accessible for many research and industry labs. Each gRNA needs to be individually designed and optimized for its DNA target. UV light was also used to activate the gRNA. To our knowledge, no approach has shown that ability to use multiple wavelengths of light to achieve multiplexing, possibly due to the reliance of UV activated photoprotecting groups. Therefore, a robust, versatile, photo-activated CRISPR-Cas9 gene editing system capable of using visible light would be highly desirable.

For these reasons we chose to explore if RNA-CLAMP could be used to control gRNA activity by site-specific cross-linking with photocleavable linkers. The mechanism of Cas9 DNA cleavage has been well studied [13, 14, 15, 16, 17]. Cas9’s endonuclease activity requires the binding of gRNA, following by the formation of the Cas9-gRNA ribonucleoprotein (RNP), which then scans through the genome until it binds to the target. The nuclease lobe of the Cas9 interacts with the protospacer adjacent motif (PAM), following by the activation of the Cas9-gRNA RNP complex. Notably, the DNA recognition lobe and the nuclease lobe of the Cas9 protein have to obtain the correct conformation to facilitate target DNA cleavage. Thus, we reasoned that if we rigidify the Cas9-gRNA RNP by cross-linking the internal loops of the gRNA, the Cas9 would lose its flexibility, resulting in loss-of-function. Subsequent cleavage of the cross-linker would remove the conformational restraint on the gRNA and activate CRISPR-Cas9 gene editing.

We applied RNA-CLAMP to enzymatically cross-link two internal stem loops within the sgRNA. Upon clamping of the sgRNA, the Cas9-sgRNA RNP completely loses its DNA cleavage activity. The subsequent photo-cleavage of the crosslinker releases the gRNA and restores gene editing activity to the wild-type level. Importantly, cross-linkers that are responsive to different wavelengths of light can be used, allowing for free selection of gene editing activation wavelength. We demonstrate multiplexed photo-activation of gene editing at two different genomic loci. The high efficiency of photo-activation enables spatiotemporal control, and we show that our technique allows gene editing within a single cell among a population of cells, which can be traced by timelapse microscopy.

## RNA-CLAMP, a site-specific RNA cross-linking technology

We previously developed a technique, RNA transglycosylation at guanosine (RNA-TAG), which enables site-specific and covalent conjugation of small molecule effectors, such as fluorophores, affinity tags, or translational regulators, onto an RNA of interest. RNA-TAG utilizes a bacterial tRNA guanine transglycosylase (TGT) to exchange a guanine nucleobase within a specific 17-nucleotide RNA stem loop structure (Tag) with a modified analog of the natural substrate preqeuosine1 (preQ1). Covalent modification is site-specific, robust, irreversible, and versatile, and RNA-TAG technology has been adapted to image cellular RNA in fixed cells, regulate mRNA translation, and study RNA-protein interactions [18–23]. We were curious whether or not RNA-TAG could be utilized to cyclize RNA targets with synthetic linkers. We imagined cyclization could be achieved by introducing two recognition sites on a single RNA, and then using a bivalent linker. The resulting internal cross-link would cyclize the RNA at the two positions where the Tag recognition site is introduced (Figure 1A). Crosslinking would likely affect the structure of the RNA, thus interfering with normal function, whereas the cleavage of the crosslinker would release the RNA, resulting in gain-of-function.

**Figure 1.**
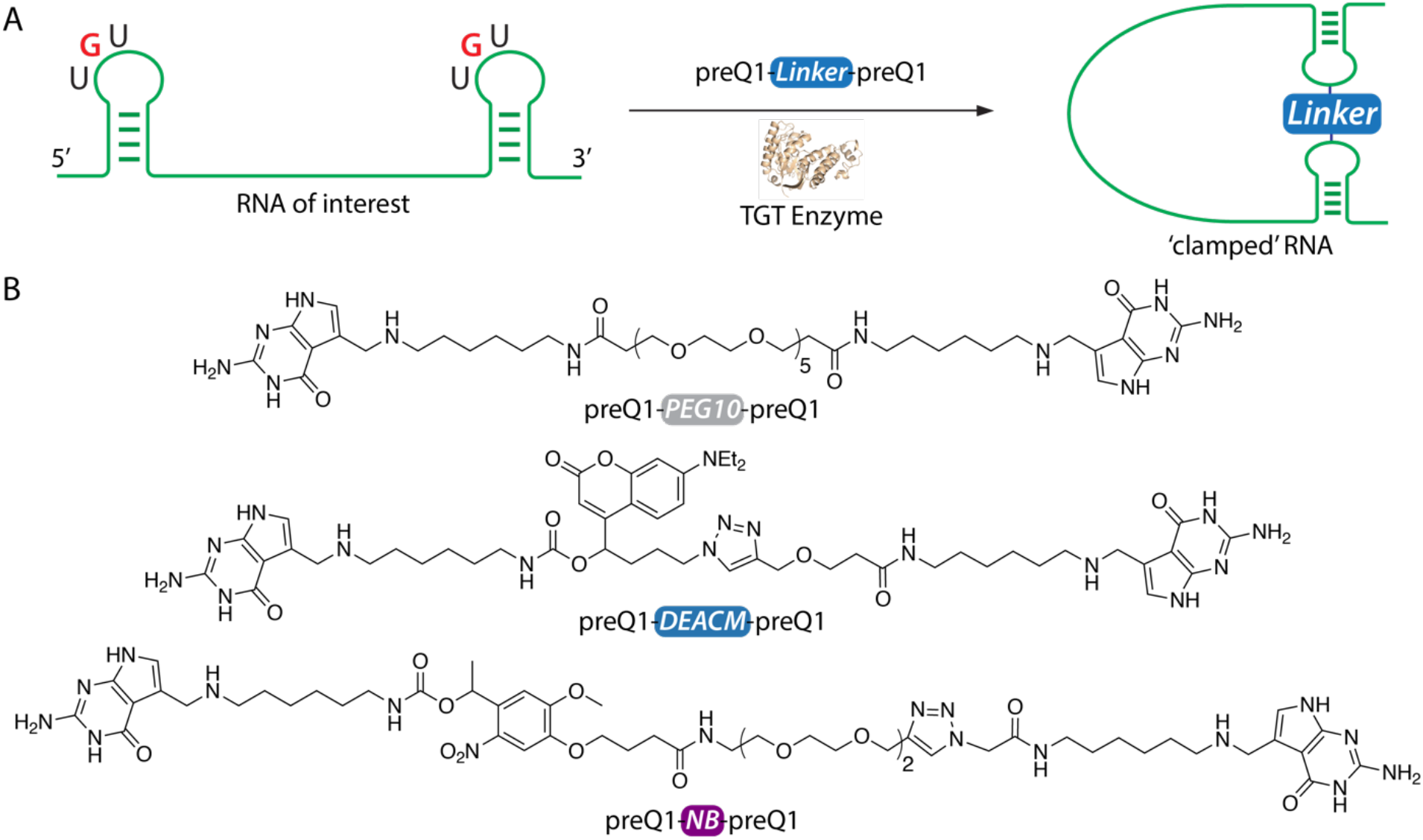
The RNA-CLAMP technique. B) Chemical structure of bivalent TGT enzymatic small-molecule substrates preQ1-PEG10-preQ1, preQ1-DEACM-preQ1 and preQ1-NB-preQ1. The preQ1-PEG10-preQ1 substrate is not photo-sensitive. The [7-(diethylamino)coumarin-4-yl]-methyl (DEACM) linker is photocleaved by irradiation with 405-456 nm light. The nitrobenzyl based (NB) linker is photo-cleaved by irradiation with 356-390 nm light.

After verifying that TGT enzymes accepts a bivalent linker (preQ1-PEG10-preQ1) and can be used to dimerize small model RNA hairpins (Figure S1), we explored if RNA-TAG can be used for RNA intramolecular cross-linking (or clamping). To enable intramolecular cross-linking, an RNA substrate needs to have two TGT recognition motifs. We designed a 74-nt long RNA molecule (RNA-1) with two Tag sequences and performed TGT enzymatic labeling using a non-cleavable preQ1-PEG10-preQ1 linker (Figure 1A). In addition to the desired clamped RNA product, byproducts were expected to form, and multiple bands were observed (Figure S2A). To distinguish between these RNA products, we designed an RNAse-H digestion assay (Figure S2B). RNAse-H specifically cleaves DNA-RNA hybrids [24, 25, 26]. Using a DNA oligo which hybridizes to the RNA, a clamped RNA should only have a single-band RNAse-H digestion pattern, whereas all byproducts should have multi-band RNAse-H digestion patterns. Denaturing TBE PAGE analysis confirmed our hypothesis (Figure S3). Therefore, we demonstrated that RNA-TAG can be used for site-specific and enzymatic cross-linking of two stem loops within an RNA of interest.

## Screening of sgRNA labeling sites

To demonstrate the applicability of RNA-CLAMP, we chose to cyclize the sgRNA of the CRISPR-Cas9 gene editing system. The sgRNA is a great candidate for our RNA-CLAMP technology since sgRNA is highly structured, as illustrated in the reported crystal structure of the sgRNA in complex with dCas9 and target DNA [13]. The sgRNA forms three stem loops as well as one tetraloop. It has been previously shown that the stem loops of the sgRNA are tolerant to mutations [13]. Since a Tag sequence is required for TGT enzymatic labeling, we first screened the tolerance of Tag sequence insertions at all of the 6 potential labeling sites of the sgRNA: the 5’-end, tetraloop, stem loop 1, stem loop 2, stem loop 3 and 3’-end (Figure S4A). Briefly, we inserted the 17-nt Tag sequence (<u>GCAGA</u>CUGUAAA<u>UCUGC</u>) at the 5’-end to form sgRNA-1 and at the 3’-end to form sgRNA-6. We swapped one of the following stem loops of the sgRNA, tetra loop, stem loop 1, stem loop 2 and stem loop 3 with the Tag sequence to form sgRNA-2, sgRNA-3, sgRNA-4 and sgRNA-5, respectively. All sgRNAs were designed to target the DYRK1A genome locus as described in a previous report. [27] To rapidly screen activity, we transiently transfected sgRNAs into HEK-293 cells that stably express Cas9 protein. 3 days after transfection, genomic DNA was harvested and the sgRNA targeted region was amplified by PCR. Sanger sequencing results were analyzed using the ICE tool to determine the INDEL rate, which is representative of the gene editing efficiency of the sgRNA [28]. We found that sgRNA-1, sgRNA-3 and sgRNA-5 had significantly reduced activity (Figure S4B). We reasoned that for sgRNA-1, the 5’ insertion of the Tag sequence might interfere with the DNA strand invasion of the sgRNA, significantly diminishing its DNA cleavage efficiency. The sgRNA-3 was not functional, likely because the RNA three-way-junction structure formed near stem loop 1 plays an important role in facilitating binding with the arginine-rich bridge helix of the Cas9 enzyme and does not tolerate sequence modification. Sequence modification on stem loop 3 (sgRNA-5) has reduced activity likely because the stem loop 3 of the sgRNA directly interacts with the Cas9 protein. However, we found that both sgRNA-2 and sgRNA-4 retained wild-type level activity. This is not surprising as the modified stem loops are solvent exposed and have been previously shown to tolerate sequence mutations [13]. Based on these findings, we constructed sgRNA-7 which has the Tag sequence inserted at both the tetraloop and stem loop 2 (Figure 2A). To our delight, the sgRNA-7 had wild-type level gene editing activity, making it the best candidate for cyclization using RNA CLAMP.

**Figure 2.**
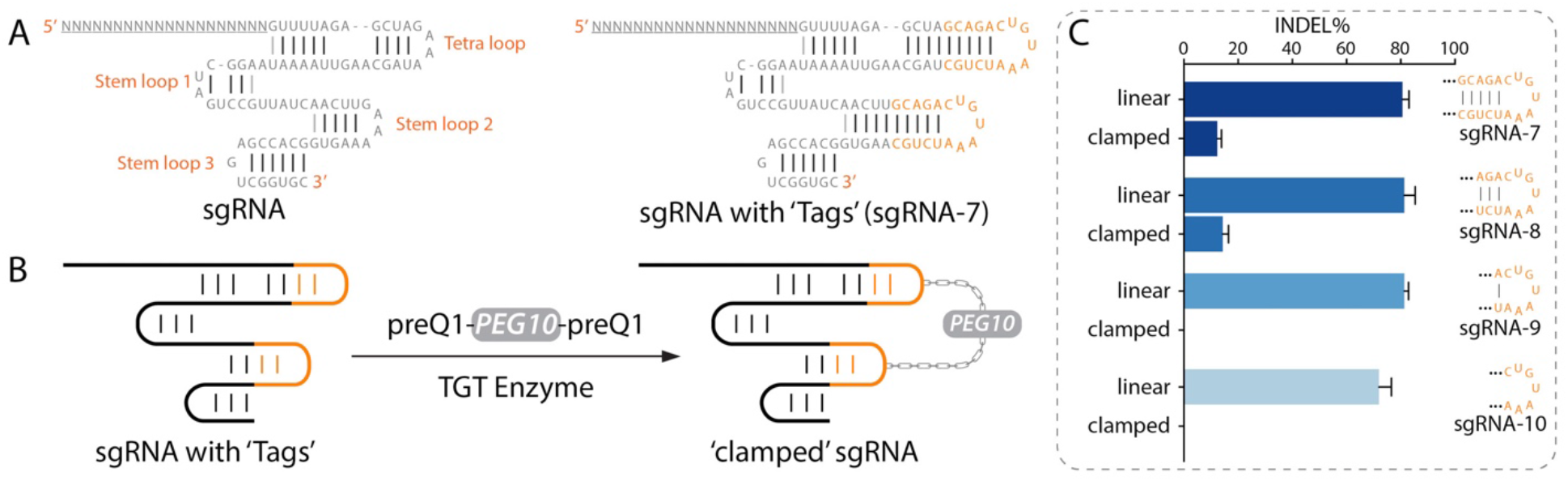
A) Sequence of the sgRNA. The sgRNA has four stem loops, which can be potentially swapped with the Tag sequence to facilitate TGT enzymatic labeling. We replaced the tetra loop and the stem loop 2 of the sgRNA to form sgRNA-7. Modification did not compromise gene editing activity. B) sgRNA-7 can be intramolecularly crosslinked using RNA-CLAMP. The ‘clamped’ sgRNA-7 completely loses its gene editing activity.

### Effect of sgRNA RNA-CLAMP on activity of CRISPR-Cas9 gene editing

We next determined the effect of intramolecular clamping on gene editing efficiency. We clamped sgRNA-7 using the non-cleavable preQ1-PEG10-preQ1 small-molecule substrate and gel purified the clamped RNA. Compared to our previous study with model RNA-1, we observed a higher yield (78.5% conversion) of the cyclized clamped sgRNA after TGT labeling. We hypothesize this is due to the sgRNA being highly structured and the orientation of the tetra loop and stem loop 2 which are facing towards the same direction. After isolating clamped sgRNA-7, we tested the gene editing activity in HEK-293 cells. The linear (unmodified) sgRNA-7 resulted in 80.7% INDEL, whereas the clamped sgRNA-7 resulted in only 12.3% INDEL. We reasoned that the flexibility of the Cas9-sgRNA complex is critical to its DNA cleavage activity. By clamping the internal loops of the sgRNA and decreasing the conformational flexibility of the RNA structure, we were able to significantly reduce the gene editing activity of the resulting CRISPR-Cas9 complex. Based on this hypothesis, we reasoned that if we shortened the distance between the two guanines that were cross-linked, further rigidifying the sgRNA, we might completely silence gene editing. TGT does not require the full 17-nt Tag sequence to label the RNA. Instead, a 7-nt (CUGUAAA) loop within a stable RNA stem structure (in this case, provided by the sgRNA backbone) is sufficient to promote TGT enzymatic labeling. Therefore, we constructed sgRNA-8, sgRNA-9 and sgRNA-10 which contained truncated stems of the Tag sequence (Figure S4A), thereby shortening the distance between the substituted guanines. We clamped sgRNA-8, sgRNA-9, sgRNA-10 and tested their gene editing activity (Figure S4C). the clamped sgRNA-8 still had 17.6% gene editing activity compared to its unmodified form. Excitingly, no detectable gene editing activity was observed with either clamped sgRNA-9 and sgRNA-10, demonstrating that the activity of the clamped sgRNA was significantly diminished by bring the two cross-linked guanine residues closer. Additionally, we found that sgRNA-9 maintained a wild-type level of gene editing activity in its unmodified form, and thus used sgRNA-9 for subsequent studies.

### Photo-activation of CRISPR-Cas9 gene editing

To enable photo-activation of CRISPR-Cas9 gene editing, we synthesized the photo-cleavable small-molecule substrate, preQ1-DEACM-preQ1. Since TGT only recognizes the preQ1 moiety, and not the linker, we had the freedom to choose from a variety of photosensitive linkers. The DEACM linker consists of a photoactive coumarin that can be cleaved by visible blue light, which minimizes photo-toxicity [29]. We clamped sgRNA-9 using the preQ1-DEACM-preQ1 substrate (79.4% yield, Figure S5A), purified the clamped sgRNA and characterized photocleavage. As shown in the denaturing PAGE analysis (Figure S5B), irradiation with a 456 nm LED light for 3 minutes completely cleaved the DEACM linker, transforming the clamped sgRNA (gel line 2 in Figure S5B) to its linear form (gel line 4 in Figure S5B). The robust enzymatic clamping and the clean photo-cleavage reaction encouraged us to pursue downstream cellular gene editing photo-activation experiments.

To test whether photo-cleavage of the crosslinker could activate the sgRNA, we delivered the linear, clamped, pre-activated sgRNA-9 into Cas9-expressing HEK293 cell line (Figure 3A). For pre-activation, clamped sgRNA-9 was irradiated with 456 nm light (LED) for 3 minutes prior to transfection. For live-cell photo-activation, sgRNA transfected cells were irradiated with 456 nm light 4 hours after transfection. 3 days later, INDEL rates were analyzed to quantify gene editing efficiency. As shown in Figure 3B, pre-activation completely restored the activity of the clamped sgRNA to the wild-type level. Live-cell photo-activation restored the clamped sgRNA-9’s activity to 84.6% compared to the wild-type. The slightly lower live-cell photo-activation might be due to degradation of the clamped sgRNA within the 4-hour transfection window. Importantly, the clamped sgRNA has zero gene editing background.

**Figure 3.**
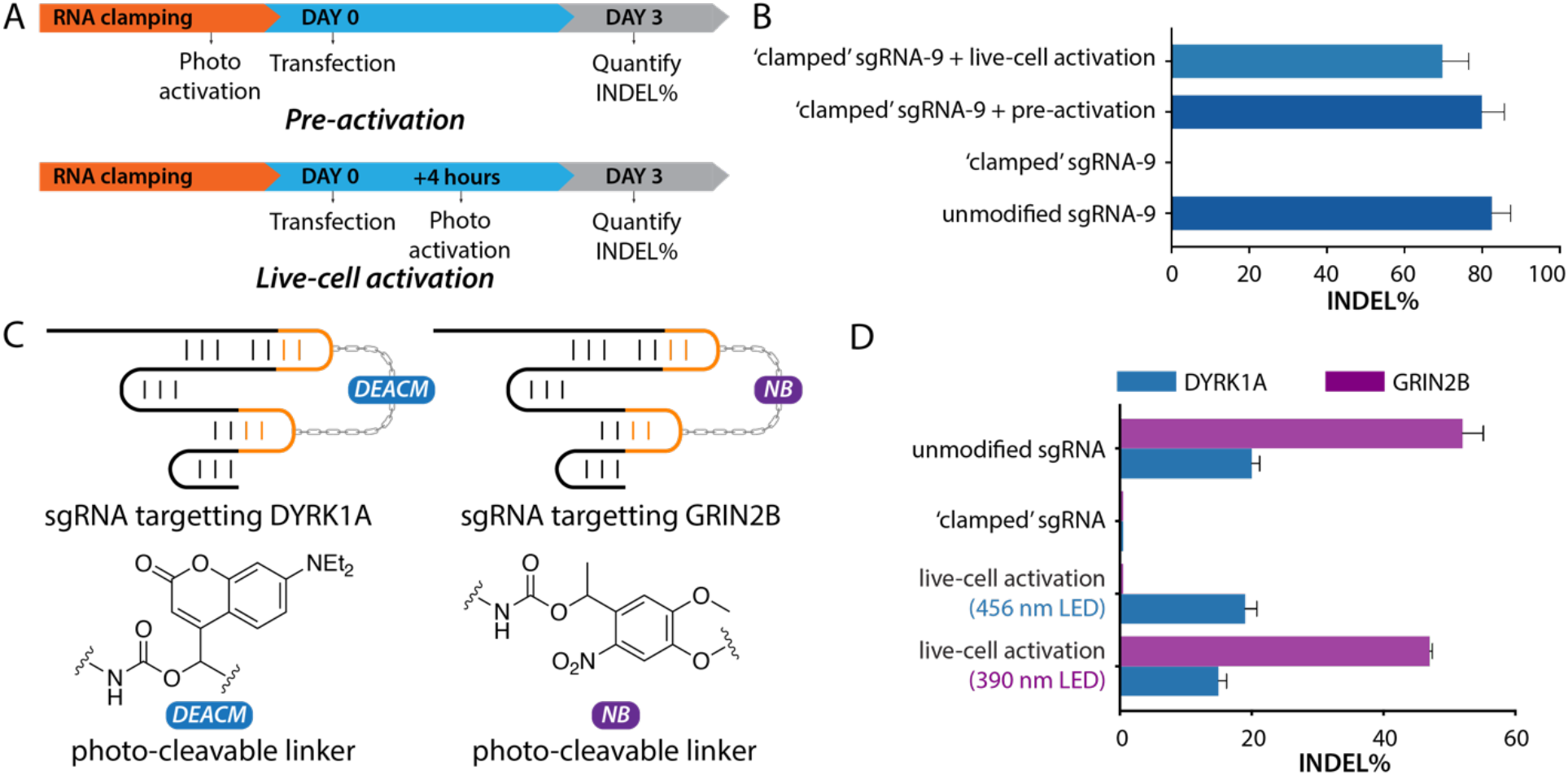
Photo-activation of CRISPR-Cas9 gene editing. A) Experimental design of pre-activation and livecell photo-activation of the ‘clamped’ sgRNA. For pre-activation, the clamped sgRNA was uncaged prior to transfection. For live-cell activation, LED light was directly applied to HEK-293 cells 4 hours after the transfection event. B) Photo-irradiation restored CRISPR-Cas9 gene editing. C) For multiplexed photoactivation of gene editing, the preQ1-DEACM-preQ1 substrate was used to clamp a sgRNA targeting the DYRK1A genomic site, and the preQ1-NB-preQ1 substrate was used to clamp a sgRNA targeting the GRIN2B genomic site. D) Multiplexed photo-activation of gene editing.

To demonstrate the spatiotemporal resolution of our CRISPR-Cas9 photo-activation technology, we adapted a surrogate reporter system which contains a series of transgenes and expresses GFP only when INDELs form at the targeted site [30]. An mCherry transgene is constitutively expressed by a CMV promoter, whereas the expression of downstream GFP genes are disrupted by an in-frame stop codon as well as a frame shift (+1 or +2 shift). Without INDEL formation, the cell will express mCherry but not GFP (Figure S7). However, when an INDEL is formed by CRISPR-Cas9 mediated gene editing and the cellular NHEJ pathway, the frame shift of the downstream GFP gene can be corrected, resulting in GFP expression (Figure S7). We inserted this series of transgenes into the genome of the HEK-293-Cas9 cells using the *Sleeping Beauty* transposition system [31]. Without gene editing, the cells only expressed mCherry while edited cells expressed both mCherry and GFP. To demonstrate spatial photo-activation of CRISPR-Cas9 gene editing, we delivered clamped sgRNA-12 targeting the sequence between the mCherry and the GFP genes into the reporter cells by transient transfection. 4 hours later, the transfection medium was replaced with complete cell-growth medium and selected cells were irradiated with a 405 nm wavelength laser. Cells were continuously imaged to observe cell growth and GFP expression. In Figure 6, the circled cell in the first image was laser irradiated for 10 seconds to activate the clamped sgRNA. 15 hours after the photo-irradiation event, cell division occurred and GFP expression was observed in one of the daughter cells, indicating INDEL formation by the CRISPR-Cas9 gene editing system. At 21.5 hours, GFP was continuously expressed in both of the daughter cells. 28.5 hours after the irradiation, one of the daughter cells undergoes mitosis, which is soon followed by mitosis in the other daughter cell. Finally, we observe a total of 4 cells expressing GFP as a result of single-cell light guided CRISPR-Cas9 gene editing and cell proliferation.

**Figure 6.**
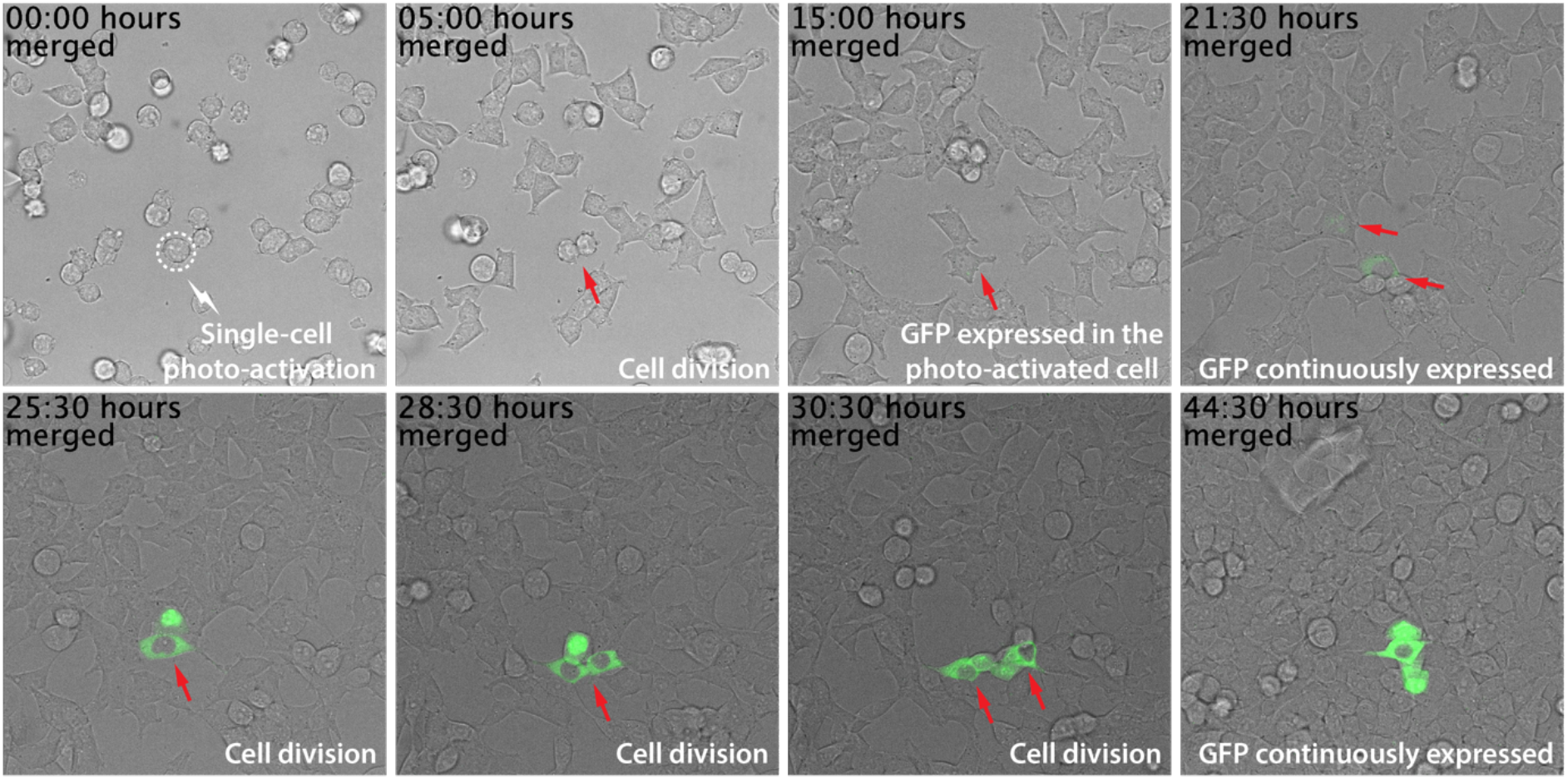
Single-cell photo-activation of CRISPR-Cas9 gene editing using a surrogate GFP reporter system.

### Multiplexed photo-activation of gene editing

One of the benefits of the CRISPR-Cas9 gene editing is its multiplexing capability. By using guide RNAs with different sequences, multiple genomic locations can be targeted at the same time. Compared to protein-centric light activation approaches, activation of the guide RNA allows the possibility for multiplexed photo-activated gene editing. The ability to spatiotemporally control editing of multiple sites could promote the study of complex gene networks with high spatiotemporal precision. One benefit of the RNA-CLAMP technology is the flexibility to choose from different photo-cleavable linkers. To demonstrate if multiplexed photo-activation of gene editing is feasible, we synthesized a nitrobenzyl (NB) based photo-cleavable linker, preQ1-NB-preQ1, which can be cleaved by irradiation with 390 nm light (Figure 1B). The NB linker requires UV light for cleavage and is not cleaved by the 456 nm light used to cleave the DEACM linker. Thus, it is possible to control photocleavage and gene editing of two genes in a wavelength specific manner. Specifically, a longer wavelength of light (456 nm) only cleaves the DEACM linker, while a shorter wavelength of light (390 nm) cleaves both linkers. We used the preQ1-NB-preQ1 to clamp sgRNA-11 which targets the GRIN2B genome locus. The sgRNA-11 has the same backbone as the sgRNA-9. Both the clamped sgRNA-9 and sgRNA-11 were transfected into Cas9-expressing HEK-293 cells. Following previously established photo-activation protocol, cells were irradiated with either 456 nm light (3 minutes) or 390 nm LED light (30 seconds) to trigger the photo-activation of gene editing (Figure 3A). Similarly, gene editing efficiency was quantified by sanger sequencing as INDEL%. As shown in Figure 3D, irradiation with the 456 nm light only triggered gene editing at the DYRK1A genome locus, while irradiation with the 390 nm wavelength of light triggered gene editing at both loci. Therefore, we demonstrated that the RNA-CLAMP photo-activated gene editing system is compatible with different wavelengths of light enabling multiplexed photo-activation of gene editing. To our knowledge, this is the first demonstration of such an approach.

### Clamped sgRNA forms Cas9-sgRNA RNPs and binds to target DNA

We asked whether clamped sgRNA maintains the capability to form the Cas9 RNP and bind to target DNA. Adapting previous reports which study the binding of sgRNA with Cas9 and the target DNA, we designed a 60 bp DNA fragment which contains the sgRNA-9 target sequence with the required NGG PAM. An in vitro gel shift assay was performed to confirm the formation of the Cas9-sgRNA RNP and its binding to the target DNA substrate (Figure S8). A non-targeting sgRNA-11 was used as the negative control. As shown in the gel shift analysis, formation of Cas9-sgRNA RNP was observed by mixing Cas9 protein with all three sgRNAs: linear sgRNA-9, clamped sgRNA-9, and linear sgRNA-11. We also observed the formation of the Cas9-sgRNA-DNA ternary complex by mixing Cas9 protein, the unmodified or ‘clamped’ sgRNA-9, and the DNA substrate. However, the Cas9-sgRNA-DNA ternary complex was not observed for sgRNA-11 (the non-targeting sgRNA), demonstrating the specificity of the CRISPR-Cas9 system. These results suggest that although the RNA clamping completely quenches the DNA cleavage activity, the clamped RNP is still able to specifically recognize and bind to its DNA target. Further work, perhaps through structural analysis of the resulting RNP complex, will be required to decipher the precise mechanism by which clamping the sgRNA prevents Cas9 function. Nonetheless, these data suggest that light activation of clamped sgRNA could trigger rapid gene editing. Ha and coworkers have previously demonstrated that photo-activation of a caged crRNA, which retains the ability to form a DNA binding Cas9 RNP, displays fast activation dynamics and enables the study of DNA damage responses and DNA repair [4b].

## Conclusion

In summary, we have developed a site-specific RNA crosslinking approach termed RNA-CLAMP. To the best of our knowledge, RNA-CLAMP is the first reported technique which allows for site-specific and enzymatic cross-linking of two internal stem loops within an RNA of interest. By incorporating a photo-cleavable linker, the clamped RNA can be released by irradiation with a user selected wavelength of light. Given the simplicity of the activation mechanism RNA-CLAMP offers great flexibility, and we demonstrated this by employing two photo-activatable crosslinkers that can be cleaved using different wavelengths of light. In principle a variety of alternative photo-cleavable linkers can be used [38]. Beyond photo-cleavable linkers, different conditionally cleavable linkers should also be applicable for clamping the RNA. For example, redox sensitive disulfide bonds, pH-sensitive linkers and enzymatically cleavable peptides [33–36]. Furthermore, the RNA-CLAMP technique can also be used to intermolecularly cross-link two RNA molecules. The versatility of our RNA-CLAMP technology will promote the development of a wide-range of biotechnologies when site-specific cross-linking of RNAs is desired.

We applied the RNA-CLAMP technique to the sgRNA of the CRISPR-Cas9 system. By exchanging the 4-nt sequence (GAAA) within the tetra loop and the stem loop 2 of the sgRNA to the 7-nt TGT recognition motif (CUGUAAA), we achieved site-specific cross-linking of the two guanine residues within these two TGT recognition motifs As a result, the Cas9-sgRNA RNP complex completely lost its DNA cleavage activity while maintaining the ability to bind to its DNA target. Live-cell photo-irradiation triggered the cleavage of the crosslinker and activated CRISPR-Cas9 gene editing. This photo-activated gene editing platform offers clean editing background as well as high cellular activation rate (84.6%). Photo-activation also offer excellent spatiotemporal precision. We were able to photo-activate CRISPR-Cas9 gene editing within a single cell among a population of cells. Furthermore, by using photo-cleavable linkers which are responsive to different wavelength of lights, we demonstrated multiplexed photo-activation of gene editing at two genomic loci. The RNA-CLAMP photo-activated CRISPR-Cas9 gene editing platform provides clean background, high activation rate, and free choice of photo-cleavable linkers to minimize photo-toxicity and enable multiplexing. We believe our approach will accelerate future development of photo-activated gene editing technologies for promoting novel biological discoveries through deciphering complex gene networks as well as leading to CRISPR based gene therapies to precisely knock in/out therapeutically relevant genes.

## Acknowledgments

We thank Xuan Zhang from Dr. Xiang-Dong Fu’s research lab at UCSD for engineering the HEK-293-Cas9 stable cell line as well as for helpful discussions on performing clamped RNA in vitro purification experiments. The project or effort depicted was sponsored by the Defense Advanced Research Projects Agency Biological Technologies Office (BTO) Safe Genes Program under Contract Number HR0011-18-2-0039 and the National Institutes of Health under grant R01 GM123285. The content of the information does not necessarily reflect the position or the policy of the government, and no official endorsement should be inferred.

