## Supplementary material for "RNA-CLAMP Enables Photo-activated Control of CRISPR-Cas9 Gene Editing by Site-specific Intramolecular Cross-linking of the sgRNA": SI

### General Materials

Reagents and instruments

Commercially available methanesulfonyl chloride, sodium azide, tetra-*n*-butylammonium fluoride in THF (1M), *N*-succinimidyl carbonate, 4-dimethylaminopyridine, *N*,*N*-diisopropylethylamine, copper(I) bromide and common organic solvents were obtained from Sigma-Aldrich. Deuterated chloroform ($CDCl_3$) was obtained from Cambridge Isotope Laboratories. All reagents obtained from commercial suppliers were used without further purification. Analytical thin-layer chromatography was performed on E. Merck silica gel 60 $F_{254}$ plates. Silica gel flash chromatography was performed using E. Merck silica gel (type 60SDS, 230-400 mesh). Solvent mixtures for chromatography are reported as v/v ratios. HPLC analysis was carried out on an Eclipse Plus C8 analytical column with *Phase A*/*Phase B* gradients [*Phase A*: $H_2O$ with 0.1% formic acid; *Phase B*: MeOH with 0.1% formic acid]. HPLC purification was carried out on a Zorbax SB-C18 semipreparative column with *Phase A*/*Phase B* gradients [*Phase A*: $H_2O$ with 0.1% formic acid; *Phase B*: MeOH with 0.1% formic acid]. Proton nuclear magnetic resonance ($^1H$ NMR) spectra were recorded on a VarianVX-500 MHz spectrometer, and were referenced relative to residual proton resonances in $CDCl_3$ (at 7.24 ppm). Chemical shifts were reported in parts per million (ppm, $\delta$) relative to tetramethylsilane (at 0.00 ppm). $^1H$ NMR splitting patterns are assigned as singlet (s), doublet (d), triplet (t), quartet (q) or pentuplet (p). All first-order splitting patterns were designated on the basis of the appearance of the multiplet. Splitting patterns that could not be readily interpreted are designated as multiplet (m) or broad (br). Electrospray Ionization-Time of Flight (ESI-TOF) spectra were obtained on an Agilent 6230 Accurate-Mass TOF mass spectrometer.

DNA oligonucleotides were purchased from Integrated DNA Technologies (Coralville, IA). Molecular biology reagents such as restriction digestion enzymes, Q5 HF-DNA polymerase, T7 RNA polymerase, nucleotide stains and competent bacterial strains were purchased from New England Biolabs (Ipswitch, MA), Promega (Madison, WI), or Life Technologies (Carlsbad, CA).

Spinning-disk confocal microscopy images were acquired on a Yokagawa spinning-disk system (Yokagawa, Japan) built around an Axio Observer Z1 motorized inverted microscope (Carl Zeiss S-3 Microscopy GmbH, Germany) with a with a 20x 0.8 NA objective to an ORCA-Flash4.0 V2 Digital CMOS camera (Hamamatsu, Japan) using ZEN Blue imaging software (Carl Zeiss Microscopy GmbH, Germany). The fluorophores were excited with diode lasers (405 nm-20 mW, 488 nm-30 mW, and 561 nm-20 mW). Images were processed using ImageJ (Fiji).

LED light sources for photo-activation

456 nm LED (50W max) light source was purchased from KESSIL.
390 nm LED (52W max) light source was purchased from KESSIL.

Reaction Buffers

TGT Storage Buffer: 25 mM HEPES, pH 7.3, 2 mM DTT, 1 mM EDTA, and 100 µM PMSF.
TGT Reaction Buffer: 100 mM HEPES, pH 7.3, 5 mM DTT, and 20 mM $MgCl_2$.
T7 Reaction Buffer: 40 mM Tris pH 7.5, 5 mM DTT, 25 mM MgCl2, 2 mM spermidine.

### Synthesis

Synthesis of preQ1-PEG10-preQ1

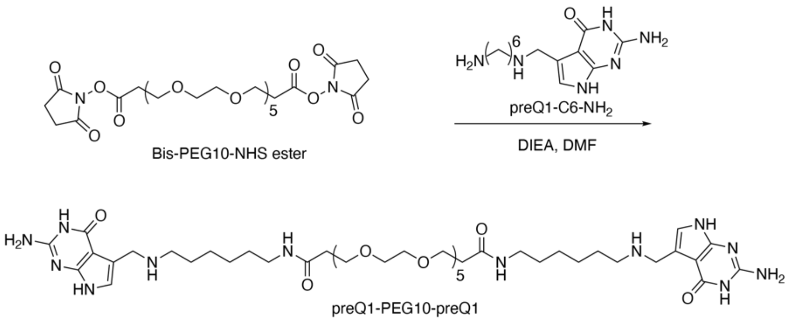

Scheme S1. Synthesis of preQ1-PEG10-preQ1.

The bis-PEG10-NHS ester was purchased from BROADPHARM (San Diego, CA). The synthesis of preQ1-C6-$NH_2$ was followed as previously described [1]. Bis-PEG10-NHS ester (6.8 mg, 9 μmol) was dissolved in 0.5 mL DMF followed by slow addition of the preQ1-C6-$NH_2$ (5.0 mg, 18 μmol in 0.5 mL DMF) and DIEA (5.8 mg, 45 μmol). The reaction solution was stirred for 1 hour at room temperature. The crude product was directly subjected to semipreparative HPLC purification, using a C18 column [gradient of $H_2O$ with 0.1% formic acid and MeOH with 0.1% formic acid 95:5 (0 min) to 5:95 (10 min to 18min)]. The semipreparative HPLC fractions containing product preQ1-PEG10-preQ1 were dried *in vacuo*, yielding 6.6 mg (6.1 μmol, 68% yield) purified product as a white residue. HRMS (M+H$^+$) calcd for $[C_{50}H_{87}N_{12}O_{14}]^+$ 1079.6459, found 1079.6460.

$^1$H NMR (500 MHz, $CD_3OD$): δ 8.55 (s, 1 H), 7.21–7.16 (m, 1 H), 6.84–6.83(m, 1 H), 4.64 (s, 1 H), 4.24–4.23 (m, 1 H), 3.74–3.72 (m, 17 H), 3.62–3.61 (m, 10 H), 3.18 (t, $J$ = 5.0 Hz, 1 H), 3.04 (t, $J$ = 5.0 Hz, 1 H), 2.42 (t, $J$ = 5.0 Hz, 1 H), 2.32–2.28 (m, 1 H), 2.17–2.15 (m, 14 H), 2.05–2.03 (m, 16 H), 1.89–1.87 (m, 16 H), 1.72 (t, $J$ = 5.0 Hz, 1 H), 1.52 (t, $J$ = 5.0 Hz, 1 H), 1.44 (t, $J$ = 5.0 Hz, 1 H), 1.35 (t, $J$ = 5.0 Hz, 1 H).

$^{13}$C NMR (126 MHz, $CD_3OD$): δ 210.14, 173.94, 162.70, 154.57, 118.15, 110.25, 99.70, 71.98, 71.51, 71.43, 71.29, 71.03, 68.87, 68.56, 67.69, 47.60, 44.89, 40.03, 37.68, 30.82, 30.74, 30.65, 30.12, 26.99, 26.65, 26.50, 26.42, 26.36.

Synthesis of preQ1-NB-preQ1

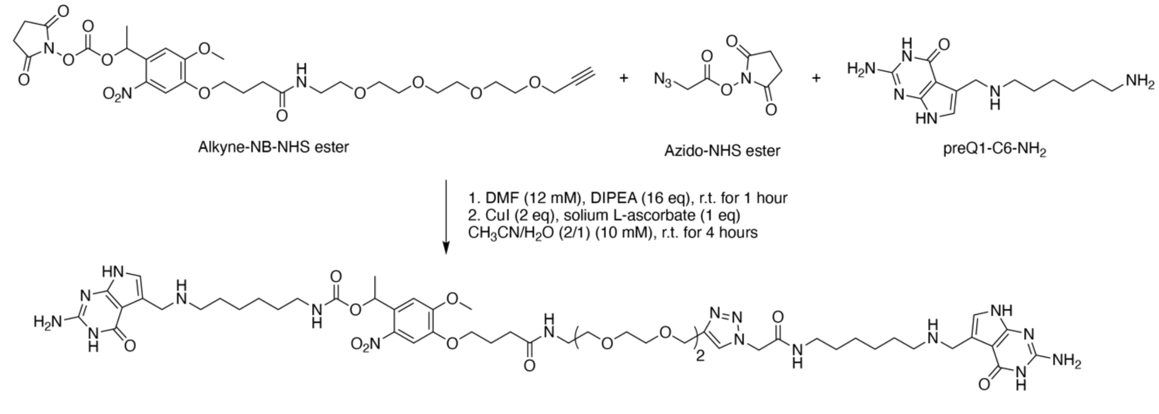

Scheme S2. Synthesis of the preQ1-NB-preQ1.

preQ1-C6-$NH_2$ was dissolved in 2 mL of anhydrous DMF. Anhydrous DIPEA (20 μL, 125 μmol) was added to the mixture. Alkyne-NB-NHS ester (8 mg, 12 μmol) and Azido-NHS ester (5 mg, 25 μmol) was added to the stirred mixture and the reaction proceeded at room temperature for 1 hour. Then the reaction solvent was removed under reduced pressure and the residue was dried *in vacuo*. Next, 1.2 mL of $CH_3CN$ : $H_2O$ (0.6 mL : 0.6 mL)，CuI (6 mg, 32 μmol), and sodium L-ascorbate (2.4 mg, 12 μmol) were added to the reaction residue.

The reaction mixture was stirred at room temperature for 4 hours. Upon completion, the reaction solvent was removed under reduced pressure and the residue was dried over high vacuum. Then $CH_2Cl_2$ was added, followed by filtration. The filtrate was concentrated by reduced pressure and the residue was purified by HPLC yielding the title compound preQ1-NB-preQ1 as a pale-yellow solid (8.9 mg, 62%). HRMS (M+H$^+$) calcd for $[C_{53}H_{80}N_{17}O_{14}]^+$ 1178.6065, found 1178.6071.

Synthesis of preQ1-alkyne

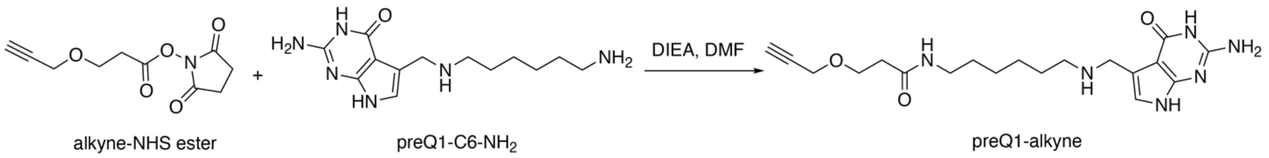

Scheme 3. Synthesis of the preQ1-alkyne

To the solution of alkyne-NHS ester (30.0 mg, 0.13 mmol, commercially available from BroadPharm, San Diego, USA; CAS#: 1174157-65-3) in DMF (3 mL) was added previously reported preQ1-C6-$NH_2$(41.9 mg, 0.13 mmol) and DIEA (38.7 mg, 0.30 mmol). The resulting solution was stirred overnight. After the substitution reaction completed, the solution was directly dried *in vacuo*. Then the mixture was purified by flash chromatography (0-50% EtOAc in hexanes) to afford the product preQ1-alkyne as pale white solid (45.9 mg, 91%). $^1$H NMR (500 MHz, Methanol-*d*4) δ 6.87 (s, 1H), 4.28 (s, 2H), 4.17 (d, *J* = 2.4 Hz, 2H), 3.79 (t, *J* = 6.1 Hz, 2H), 3.38 (s, 1H), 3.24 – 3.18 (m, 2H), 3.08 (t, *J* = 7.5 Hz, 2H), 2.89 (t, *J* = 2.4 Hz, 1H), 2.46 (t, *J* = 6.1 Hz, 2H), 1.76 (p, *J* = 7.5 Hz, 2H), 1.56 (p, *J* = 7.1 Hz, 2H), 1.51 – 1.36 (m, 4H). $^{13}$C NMR (126 MHz, Methanol-*d*4, δ): δ 172.09, 161.06, 152.92, 152.39, 117.49, 108.24, 98.06, 78.87, 74.52, 74.25, 65.43, 57.27, 45.99, 43.22, 38.44, 35.92, 28.48, 25.67, 15.71. HRMS (M+H$^+$) calcd for $[C_{19}H_{29}N_6O_3]^+$ 389.2298, found 389.2296.

Synthesis of preQ1-DEACM-preQ1

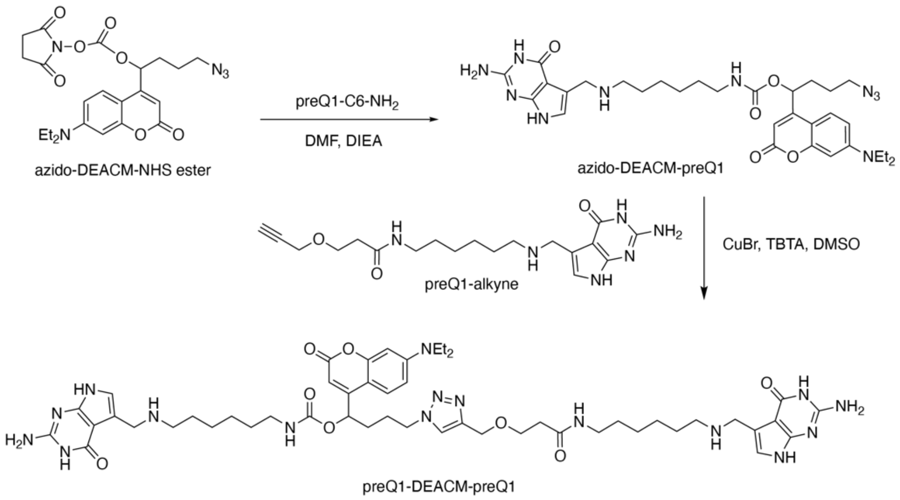

Scheme 4. Synthesis of preQ1-DEACM-preQ1

The preparation of the azido-DEACM-HNS ester is described in previous report [5]. preQ1-C6-$NH_2$ (8.8 mg, 0.032 mmol) and DIEA (7.1 mg, 0.055 mmol) in DMF (0.5 mL) was added to azido-DEACM-NHS ester (5.0 mg, 0.011 mmol) in DMF (0.5 mL). The resulting solution was stirred for 1 hour. After the substitution completed, the solution was directly subjected to semipreparative HPLC purification, using a C18 column [gradient of $H_2O$ with 0.1% formic acid and MeOH with 0.1% formic acid 95:5 (0 min) to 5:95 (10 min to 18min)]. Shielded from light, the fractions containing the product azido-DEACM-preQ1 [confirmed by low resolution MS, LRMS ($M+H^+$) calcd for $[C_{31}H_{43}N_{10}O_5]^+$ 635.3, found 635.3] from semipreparative HPLC purification were dried *in vacuo* with ice bath. Otherwise, the product azido-DEACM-preQ1 is prone to decompose. After drying *in vacuo* with ice bath, there is a little amount of water left in the vial, due to the low temperature, which was directly applied to the next step without further processing.

Shielded from light, CuBr [0.8 mg, 0.0055 mmol in DMSO : $H_2O$ (0.3 ml : 0.3 mL)] was added to the vial, followed by the addition of preQ1-alkyne (4.3 mg, 0.011 mmol) in DMSO (0.4 mL). The resulting solution was stirred for about 2 hours. After the click reaction completed, the solution was directly subjected to semipreparative HPLC purification, using a C18 column [gradient of $H_2O$ with 0.1% formic acid and MeOH with 0.1% formic acid 95:5 (0 min) to 5:95 (10 min to 18min)]. Shielded from light, the fractions containing the product preQ1-DEACM-preQ1 from semipreparative HPLC purification were dried *in vacuo* at room temperature. After drying *in vacuo* with ice bath, there was a little amount of water left in the vial, due to the low temperature. Then, a solvent mixture of $H_2O$ : MeCN (0.3 ml : 0.3 mL) was added to the vial. After freezing lyophilization, preQ1-DEACM-preQ1 (0.5 mg, 4.4 % for 2 steps) was obtained, confirmed by HRMS. HRMS ($M+2H^+$) calcd for $[C_{50}H_{72}N_{16}O_8]^+$ 512.2850, found 512.2854.

### RNA labeling using the RNA-CLAMP technology

Dimerization of the 17-nt Tag RNA oligo by RNA-CLAMP

17-nt RNA Tag oligo (GCAGACUGUAAAUCUGC) was purchased from IDT. TGT labeling reaction was assembled with the following components in 1X TGT reaction buffer: 10 µM of 17-nt RNA oligo, 5 µM of TGT enzyme, 5 µM of small-molecule substrate (preQ1-PEG10-preQ1, Figure S1A), and 5 mM DTT. The reaction mixture was incubated at 37 °C for 4 hours. 1 µL of proteinase K was added into reaction mixture and incubated at 37°C for 30 minutes to terminate the labeling reaction. The reaction was analyzed on 18% denaturing PAGE (Figure S1B). In Figure S1B, we can see that after the TGT labeling reaction, two new RNA products were produced. The top RNA band is the dimerized 17-nt Tag oligo. The middle band is the singly labeled Tag oligo. This experiment demonstrated the capability of the TGT enzyme to form a covalent bond between two RNA hairpins and builds the foundation for our RNA-CLAMP technology.

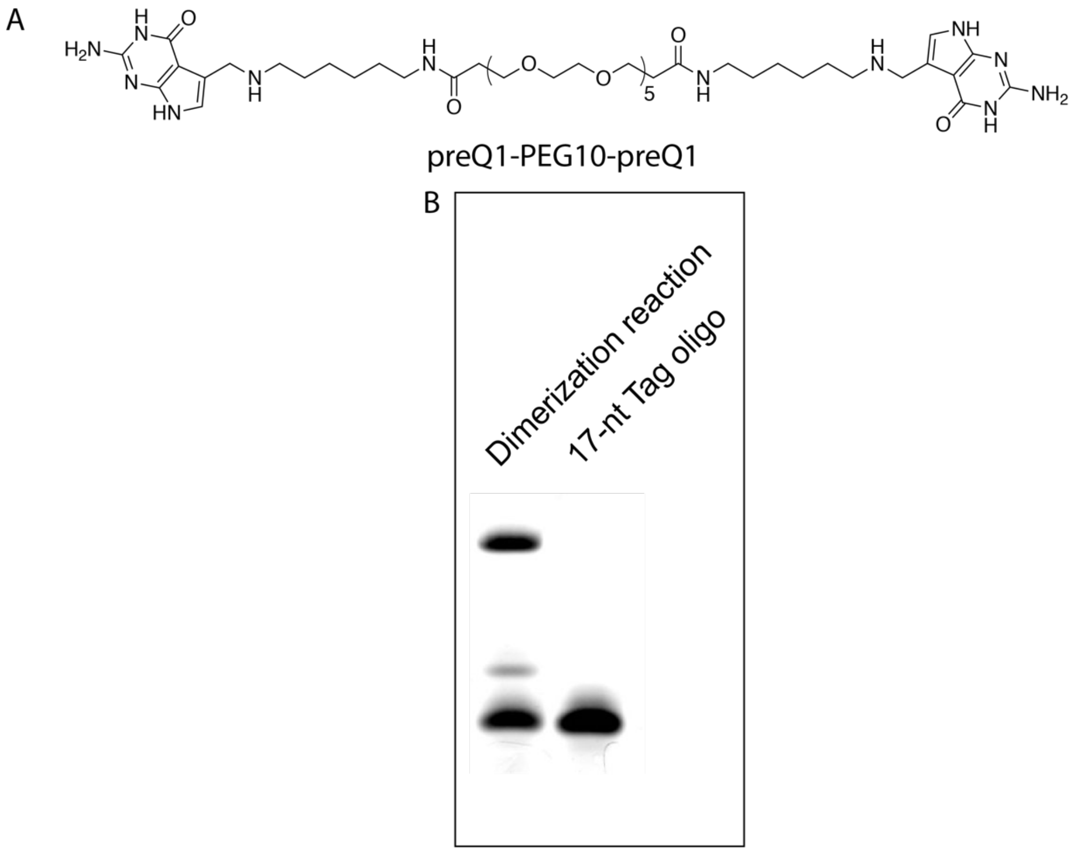

Figure S1. 18% denaturing PAGE analysis of in vitro TGT labeling of the 17-nt RNA oligo using the preQ1-PEG10-preQ1 small-molecule substrate. Gel was stained with 1X GelRed.

Intramolecular cross-link of the RNA

Sequence of the RNA-1 (the Tag sequences are underlined)
GCAGACTGTAAATCTGCCACTAGTAACGGCCGCCAGTGTGCTGGAATTCTGCAGATAGCAGACTGTAAAT
CTGC
The in vitro transcription reaction was performed following previous established protocols [1]. PAGE gel purified RNA-1 was subjected to TGT labeling reaction using the preQ1-PEG10-preQ1 small-molecule substrate. TGT labeling reaction was assembled with the following components in 1X TGT reaction buffer: 1 μM of RNA-1, 2 μM of TGT enzyme, 1 μM of small-molecule substrate (preQ1-PEG10-preQ1, Figure S1A), 1 unit/μL of Super RNase Inhibitor (Invitrogen, catalog number AM2694), and 5 mM DTT. The reaction mixture was incubated at 37 °C for 4 hours. Next, 1 μL of proteinase K was added into the reaction mixture and incubated at 37°C for 30 minutes to terminate the labeling reaction and get rid of potential RNAse contamination. Note: adding the small-molecule substrate last should promote higher 'clamping' conversion. The reason is that the TGT enzyme forms a covalent bound with the RNA substrate, resulting in a TGT-RNA intermediate. Adding the small-molecule substrate last ensured that two TGT molecules were covalently tethered to the RNA. In this manner, intermolecular cross-linking of RNA can be reduced. The crude TGT labeling products were analyzed by 7% denaturing TBE PAGE (Figure S2A). As shown in Figure S2A, we observed multiple RNA products. First, we observed unlabeled RNA starting material, shown as the bottom band with the same migration distance as Line 1. We also observed RNA degradation shown as smear bands below the RNA starting material. We hypothesize that, apart from the desired 'clamped' RNA product, the

TGT labeling reaction should at least generate two kinds of by-products: by-product 1 was generated by attaching one small-molecule substrate at each Tag sequence without forming intermolecular or intramolecular cross-linking; by-product 2 was formed through intermolecular cross-linking, which were shown as multiple top bands in Line 2.

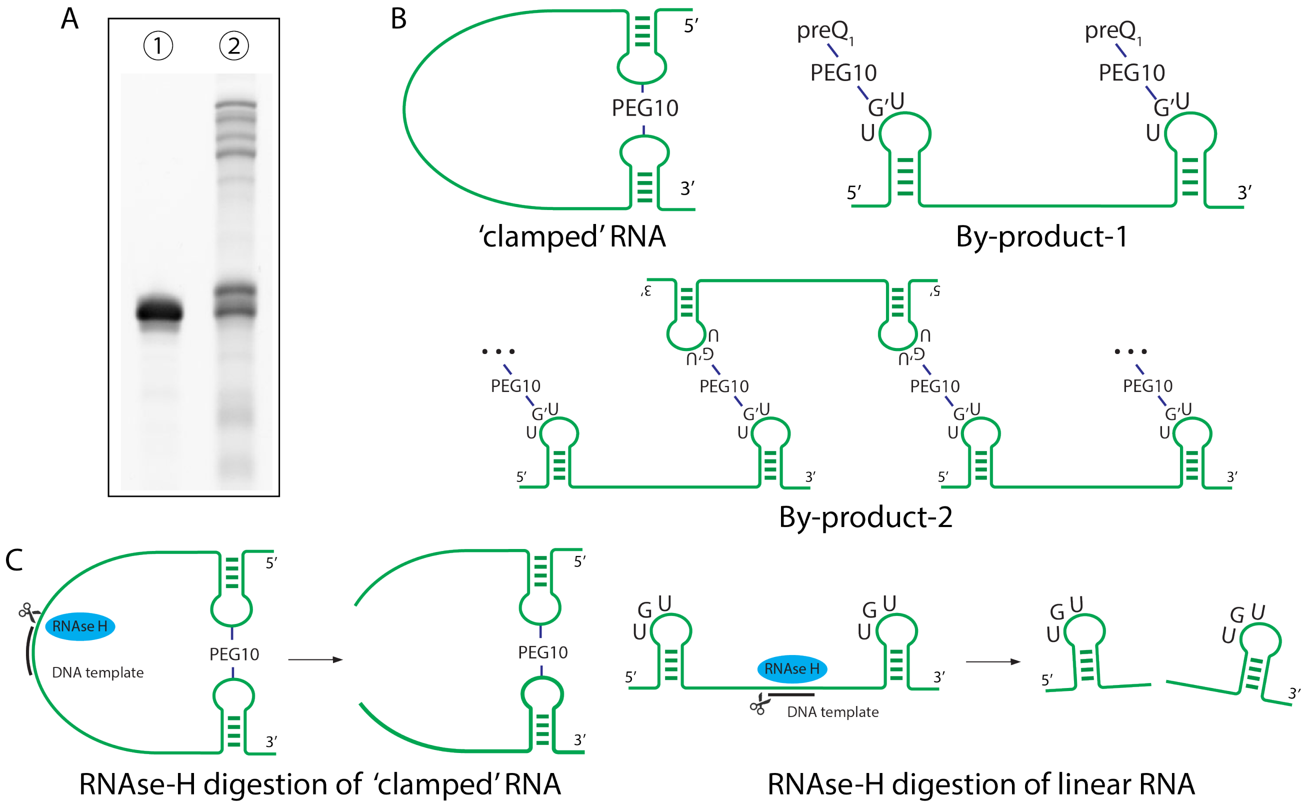

Figure S2. 7% denaturing TBE PAGE analysis of the RNA clamping reaction. A) Line1: RNA-1 starting material. The previously purified RNA starting material (RNA-1) was presented as a major single band. Line2: multiple RNA products were observed after the TGT labeling reaction.

To distinguish between multiple RNA products after the TGT reaction, we designed an RNAse-H digestion assay (Figure S2C). RNAse-H only digests RNA-DNA hybrids. We designed a DNA oligo which was complementary to the internal sequence of RNA-1 and use RNAse-H to specifically cut the internal sequence of RNA-1. As illustrated in Figure S2C, after the RNAse-H digestion (DNA oligo used for the RNAse-H digestion assay on RNA-1: CCAGCACACTGGCGGCCG), linear RNA products would generate two RNA fragments whereas the 'clamped' RNA product would only generate one RNA digestion product. We separated the product bands observed on Figure S2A (Line 2) and ran each RNA product through the RNAse-H digestion assay (Figure S3). As expected, both the RNA byproduct-1 and byproduct-2 generated two or multiple bands after the RNAse-H digestion assay. Only the 'clamped' RNA product generated one single band after the RNAse-H digestion assay. Therefore, we verified that TGT can successfully form an intramolecular cross-link (or 'clamp') on the RNA of interest using the preQ1-PEG10-preQ1 small-molecule substrate.

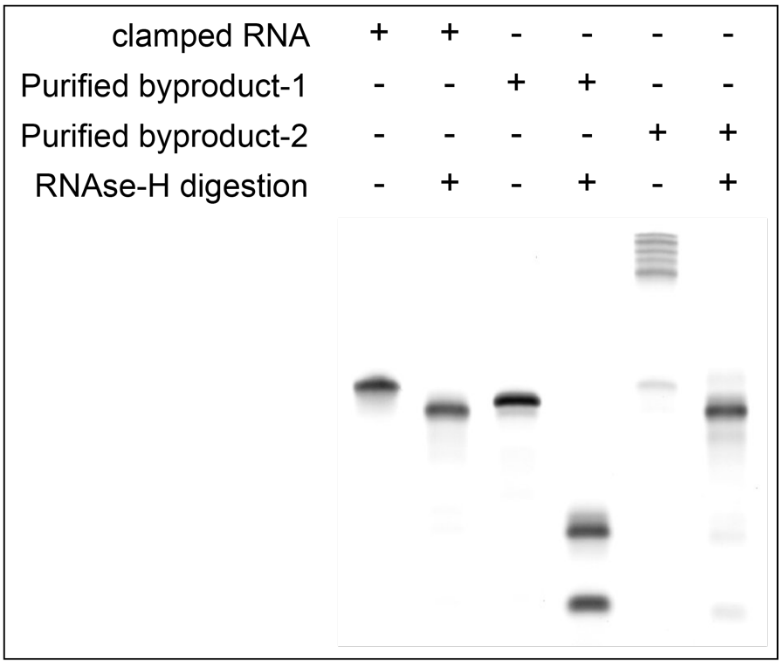

Figure S3. RNAse-H digestion assay on the RNA products after TGT clamping reaction.

### Synthesis of the single-guide RNA (sgRNA)

In vitro transcription reaction template

In vitro transcription template was purchased from IDT as ssDNA and used directly for in vitro transcription reactions.

In vitro transcription template (ssDNA, 20 nt targeting sequence was shown as $N^{20}$):

| | |
|---|---|
| DYRK1A targeting sequence | GCTGCTGGCCTTCAGATGGC |
| GRIN2B targeting sequence | GGAGAACAGCACTCCGCTCT |

wt-sgRNA:

CTAATACGACTCACTATANNNNNNNNNNNNNNNNNNNNGTTTTAGAGCTAGAAATAGCAAGTTAAAATAA
GGCTAGTCCGTTATCAACTTGAAAAAGTGGCACCGAGTCGGTGCTTTTTT

sgRNA-1:

CTAATACGACTCACTATAGCAGACTGTAAATCTGCTTTTNNNNNNNNNNNNNNNNNNNNGTTTTAGAGCT
AGAAATAGCAAGTTAAAATAAGGCTAGTCCGTTATCAACTTGAAAAAGTGGCACCGAGTCGGTGCTTTTTT

sgRNA-2:

CTAATACGACTCACTATANNNNNNNNNNNNNNNNNNNNGTTTTAGAGCTAGCAGACTGTAAATCTGCTAG
CAAGTTAAAATAAGGCTAGTCCGTTATCAACTTGAAAAAGTGGCACCGAGTCGGTGCTTTTTT

sgRNA-3:

CTAATACGACTCACTATANNNNNNNNNNNNNNNNNNNNGTTTTAGAGCTAGAAATAGCAAGTTAAAATAA
GGCGCAGACTGTAAATCTGCGTCCGTTATCAACTTGAAAAAGTGGCACCGAGTCGGTGCTTTTTTT

sgRNA-4:

CTAATACGACTCACTATANNNNNNNNNNNNNNNNNNNNGTTTTAGAGCTAGAAATAGCAAGTTAAAATAA
GGCTAGTCCGTTATCAACTTGCAGACTGTAAATCTGCAAGTGGCACCGAGTCGGTGCTTTTTT

sgRNA-5:
CTAATACGACTCACTATANNNNNNNNNNNNNNNNNNNNGTTTTAGAGCTAGAAATAGCAAGTTAAAATAA
GGCTAGTCCGTTATCAACTTGAAAAAGTGGCACCGAGCAGACTGTAAATCTGCTCGGTGCTTTTTT
sgRNA-6:
CTAATACGACTCACTATANNNNNNNNNNNNNNNNNNNNGTTTTAGAGCTAGAAATAGCAAGTTAAAATAA
GGCTAGTCCGTTATCAACTTGAAAAAGTGGCACCGAGTCGGTGCTTATTGCAGACTGTAAATCTGCTTTTT
T
sgRNA-7:
CTAATACGACTCACTATANNNNNNNNNNNNNNNNNNNNGTTTTAGAGCTAGCAGACTGTAAATCTGCTAG
CAAGTTAAAATAAGGCTAGTCCGTTATCAACTTGCAGACTGTAAATCTGCAAGTGGCACCGAGTCGGTGC
TTTTTT
sgRNA-8:
CTAATACGACTCACTATANNNNNNNNNNNNNNNNNNNNGTTTTAGAGCTAAGACTGTAAATCTTAGCAAG
TTAAAATAAGGCTAGTCCGTTATCAACTTAGACTGTAAATCTAAGTGGCACCGAGTCGGTGCTTTTTT
sgRNA-9:
CTAATACGACTCACTATANNNNNNNNNNNNNNNNNNNNGTTTTAGAGCTAACTGTAAATTAGCAAGTTAA
AATAAGGCTAGTCCGTTATCAACTTACTGTAAATAAGTGGCACCGAGTCGGTGCTTTTTT
sgRNA-10:
CTAATACGACTCACTATANNNNNNNNNNNNNNNNNNNNGTTTTAGAGCTACTGTAAATAGCAAGTTAAAA
TAAGGCTAGTCCGTTATCAACTTCTGTAAAAAGTGGCACCGAGTCGGTGCTTTTTT

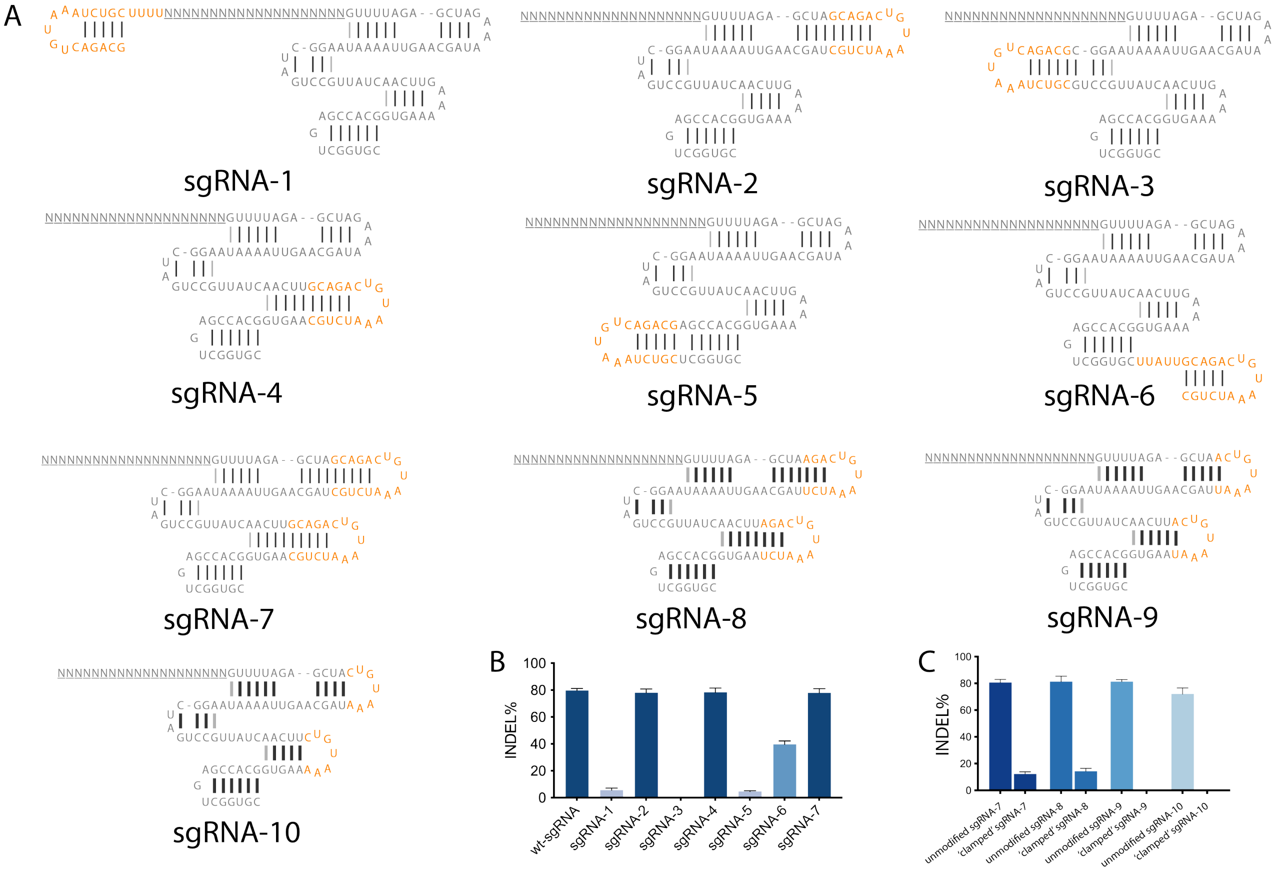

Figure S4. Sequences of sgRNA-1 to sgRNA-7 and their gene editing activities.

In vitro transcription reaction

Each IVT reaction was set up with 50 nM of linearized DNA template, 5 mM of each ATP, CTP, UTP, 9 mM of GTP (NEB, Ipswitch, MA), 0.004 unit/μL of thermostable inorganic pyrophosphatase (NEB, Ipswitch, MA), 0.25 μg/μL T7 RNA polymerase, 0.05% Triton X-100 (Sigma, St. Louis, MO) and 1 unit/μL RNase Inhibitor, Murine (NEB, Ipswitch, MA). The IVT reaction was carried out at 37 °C for 4 hours to allow for sufficient RNA synthesis. To remove DNA template, 2 μL of 100 mM $CaCl_2$ and 20 units of Turbo DNase (Life Technologies, Carlsbad, CA) were added to the mixture and incubated at 37 °C for 1 hour. The mixture was then centrifuged at 10,000 RCF for 5 minutes at room temperature to pellet any remaining magnesium pyrophosphate. The supernatant was resuspended in 50 μL RNase free water, followed by 12% denaturing PAGE purification. Purified IVT products were stored at -80°C until used.

### Intramolecular cross-link of the sgRNA by RNA-CLAMP

RNA-CLAMP using preQ1-DEACM-preQ1

To 'clamp' the sgRNA-9, 1 μM of sgRNA transcript, 1 μM of TGT enzyme, 1 μM of small-molecule substrate, 5 mM of DTT, 1 unit/μL of RNAse Inhibitor was assembled in 1X TGT buffer. The reaction mixture was incubated at 37°C for 4 hours, followed by the addition of 1 μL of proteinase K. The crude labeling products were analyzed by 12% denaturing TBE-PAGE (Figure S5A). As shown in Figure S5A, the conversion rate to the 'clamped' sgRNA was 79.4%, which is much higher than the conversion rate of the RNA-1. We reasoned that this was due to the stable ternary structure of the sgRNA. The first Tag sequence was located at the tetra-loop of the sgRNA and the second Tag sequence was located at the stem loop 2 of the sgRNA. These two loops are well-structured and close to each other, promoting higher intramolecular cross-linking conversion. To get rid of undesired RNA product and reduce background gene editing activity, 'clamped' sgRNA-9 was purified by denaturing TBE-PAGE. The purified 'clamped' sgRNA-9 was shown as a single band on a 12% denaturing PAGE gel (Figure S5B). To uncage the 'clamped' sgRNA-9, a 456 nm LED light (50W max) was used to irradiate the 'clamped' sgRNA in water for 3 minutes. LED irradiation completely photo-cleaved the DEACM linker, transforming the 'clamped' sgRNA to its linear form (Figure S5B).

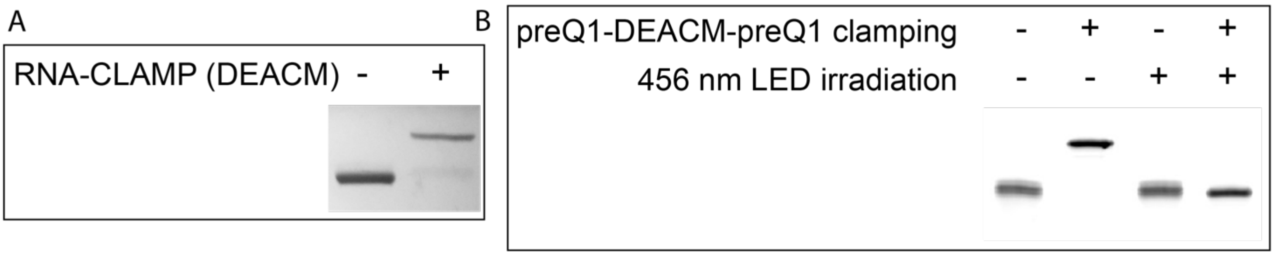

Figure S5. 'Clamping' of the sgRNA-9 using preQ1-DEACM-preQ1 substrate. A) 12% PAGE analysis of the RNA-CLAMP reaction on sgRNA-9 using the preQ1-DEACM-preQ1 small-molecule substrate. B) In vitro photo-cleavage assay of the unmodified and the 'clamped' sgRNA-9.

RNA-CLAMP using preQ1-NB-preQ1

Similarly, we performed RNA-CLAMP reaction on sgRNA-11 using the preQ1-NB-preQ1 small molecule substrate. As shown in Figure S6, the 'clamped' sgRNA-11 can be completely photo-uncaged by irradiation with a 390 nm LED light (52W max) for 30 seconds. As a control experiment, we irradiated the 'clamped' sgRNA-11 using the 456 nm LED light for 3 minutes. Only minimal photo-cleavage of the NB linker was observed (0.09%), demonstrating that the NB linker can be efficiently cleaved by a 390 nm LED, but not a 456 nm LED.

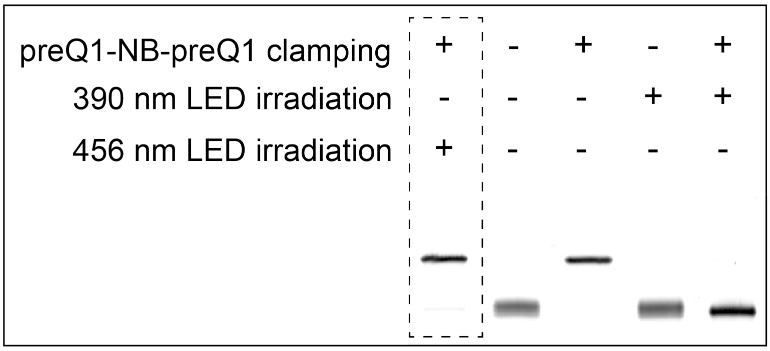

Figure S6. 'Clamping' of the sgRNA-11 using preQ1-NB-preQ1 substrate and the in vitro photo-uncaging.

### CRISPR-Cas9 gene editing

Gene editing in HEK-293-Cas9 cell line

Gene editing experiments were performed following previously reported protocols [2]. For gene editing experiments using HEK-293-Cas9 cell line, 250 ng sgRNA and 0.5 µL Lipofectamine RNAiMAX were used to form the Lipo-RNA complex. The formed complex was added with 150 µL DMEM (10% FBS) containing 40K HEK-293-Cas9 cells. The transfection was performed in a 96-well plate format. 24 hours after transfection, cell medium was changed to complete DMEM medium. At day 4, genomic DNA was extracted using 80 µL QuickExtract™ DNA Extraction Solution (Lucigen, catalog number: QE09050). Following the manufacturer's protocol, the extraction solution was heated at 65°C for 15 minutes and then at 95°C for another 15 minutes. The resulting solution was diluted using water at a 1:10 ratio. The diluted product was directly used as a template for PCR amplification of the genomic region. For PCR amplification of the genomic region, Q5 HOTSTART HIFI DNA polymerase 2X master mix was used following the manufacturer's protocol without further optimization. The primers used for amplifying the DYRK1A and GRIN2B genome locus are listed below.

| | |
|---|---|
| DYRK1A_fwd | GGAGCTGGTCTGTTGGAGAA |
| DYRK1A_rvs | TCCCAATCCATAATCCCACGTT |
| GRIN2B_fwd | CAGGAGGGCCAGGAGATTTG |
| GRIN2B_rvs | TGAAATCGAGGATCTGGGCG |

Crude PCR products were column purified using a Zymo DNA purification kit. The purified PCR amplicon was analyzed by Sanger sequencing to estimate INDEL% using the ICE tool [3]. The wt-sgRNA gene editing INDEL% was usually around 70-80% following this protocol.

Gene editing in HEK-293 cell line

For gene editing experiments performed using a HEK-293 wild-type cell line, mRNA encoding the Cas9 protein (Trilink, catalog number L-7206-20) was transfected into HEK-293 cells one day prior to sgRNA transfection. For mRNA transfection, Lipo-RNA complex was formed by using 500 ng mRNA, 0.9 µL Lipofectamine RNAiMAX in 50 µL total volume, combined with 150 µL DMEM complete medium containing 40K HEK-293 cells. At day 2, sgRNA transfection was performed following the protocol described in the previous section. The wt-sgRNA gene editing INDEL% was usually around 40-50% following this protocol.

### Surrogate EGFP reporter

The surrogate EGFP reported was adopted from a previously reported method [4]. In brief, an mCherry transgene is constitutively expressed by a CMV promoter, whereas the expression of downstream GFP genes are disrupted by an in-frame stop codon as well as a frame shift (+1 or +2 shift). Without INDEL formation, the cell will express mCherry but not GFP. However, when an INDEL is formed by CRISPR-Cas9 mediated gene editing and the cellular NHEJ pathway, the frame shift of the downstream GFP gene can be corrected, resulting in GFP expression.

Sequence of the EGFP surrogate reporter

ATGGTGAGCAAGGGCGAGGAGGATAACATGGCCATCATCAAGGAGTTCATGCGCTTCAAGGTGCACATG
GAGGGCTCCGTGAACGGCCACGAGTTCGAGATCGAGGGCGAGGGCGAGGGCCGCCCCTACGAGGGCA
CCCAGACCGCCAAGCTGAAGGTGACCAAGGGTGGCCCCCTGCCCTTCGCCTGGGACATCCTGTCCCCT
CAGTTCATGTACGGCTCCAAGGCCTACGTGAAGCACCCCGCCGACATCCCCGACTACTTGAAGCTGTCC
TTCCCCGAGGGCTTCAAGTGGGAGCGCGTGATGAACTTCGAGGACGGCGGCGTGGTGACCGTGACCCA
GGACTCCTCCCTGCAAGACGGCGAGTTCATCTACAAGGTGAAGCTGCGCGGCACCAACTTCCCCTCCGA
CGGCCCCGTAATGCAGAAGAAGACGATGGGCTGGGAGGCCTCCTCCGAGCGGATGTACCCCGAGGAC
GGCGCCCTGAAGGGCGAGATCAAGCAGAGGCTGAAGCTGAAGGACGGCGGCCACTACGACGCTGAGG
TCAAGACCACCTACAAGGCCAAGAAGCCCGTGCAGCTGCCCGGCGCCTACAACGTGAACATCAAGTTGG
ACATCACCTCCCACAACGAGGACTACACCATCGTGGAACAGTACGAACGCGCCGAGGGCCGCCACTCC
ACCGGCGGCATGGACGAGCTGTACAAGgaGCTAGCATT**<u>GTTCAGGGCTTGACCAACAC</u>**TGGGTCCCTGC
AGGatccagtgaGCAAGGGCGAGGAGCTGTTCACCGGGGTGGTGCCCATCCTGGTCGAGCTGGACGGCGA
CGTAAACGGCCACAAGTTCAGCGTGTCCGGCGAGGGCGAGGGCGATGCCACCTACGGCAAGCTGACCC
TGAAGTTCATCTGCACCACCGGCAAGCTGCCCGTGCCCTGGCCCACCCTCGTGACCACCCTGACCTACG
GCGTGCAGTGCTTCAGCCGCTACCCCGACCACATGAAGCAGCACGACTTCTTCAAGTCCGCCATGCCCG
AAGGCTACGTCCAGGAGCGCACCATCTTCTTCAAGGACGACGGCAACTACAAGACCCGCGCCGAGGTG
AAGTTCGAGGGCGACACCCTGGTGAACCGCATCGAGCTGAAGGGCATCGACTTCAAGGAGGACGGCAA
CATCCTGGGGCACAAGCTGGAGTACAACTACAACAGCCACAACGTCTATATCATGGCCGACAAGCAGAA
GAACGGCATCAAGGTGAACTTCAAGATCCGCCACAACATCGAGGACGGCAGCGTGCAGCTCGCCGACC
ACTACCAGCAGAACACCCCCATCGGCGACGGCCCCGTGCTGCTGCCCGACAACCACTACCTGAGCACC
CAGTCCGCCCTGAGCAAAGACCCCAACGAGAAGCGCGATCACATGGTCCTGCTGGAGTTCGTGACCGC
CGCCGGGATCACTCTCGGCATGGACGAGCTGTACAAGgttataacatgggtaacctctacgtgagcaagggcgaggAGCT
GTTCACCGGGGTGGTGCCCATCCTGGTCGAGCTGGACGGCGACGTAAACGGCCACAAGTTCAGCGTGT
CCGGCGAGGGCGAGGGCGATGCCACCTACGGCAAGCTGACCCTGAAGTTCATCTGCACCACCGGCAAG
CTGCCCGTGCCCTGGCCCACCCTCGTGACCACCCTGACCTACGGCGTGCAGTGCTTCAGCCGCTACCC
CGACCACATGAAGCAGCACGACTTCTTCAAGTCCGCCATGCCCGAAGGCTACGTCCAGGAGCGCACCAT

CTTCTTCAAGGACGACGGCAACTACAAGACCCGCGCCGAGGTGAAGTTCGAGGGCGACACCCTGGTGA
ACCGCATCGAGCTGAAGGGCATCGACTTCAAGGAGGACGGCAACATCCTGGGGCACAAGCTGGAGTAC
AACTACAACAGCCACAACGTCTATATCATGGCCGACAAGCAGAAGAACGGCATCAAGGTGAACTTCAAGA
TCCGCCACAACATCGAGGACGGCAGCGTGCAGCTCGCCGACCACTACCAGCAGAACACCCCCATCGGC
GACGGCCCCGTGCTGCTGCCCGACAACCACTACCTGAGCACCCAGTCCGCCCTGAGCAAAGACCCCAA
CGAGAAGCGCGATCACATGGTCCTGCTGGAGTTCGTGACCGCCGCCGGGATCACTCTCGGCATGGACG
AGCTGTACAAGTAA

Activity of the surrogate EGFP reporter in HEK-293 cells

To test the surrogate reporter in HEK-293 cells. The preQ1-DEACM-preQ1 ‘clamped’ sgRNA-12 targeting the sequence between the mCherry and the GFP genes was introduced into the surrogate EGFP reporter cells by transient transfection (sgRNA targeting sequence:  GTTCAGGGCTTGACCAACAC). 4 hours later, the transfection medium was replaced with complete cell-growth medium and cells were irradiated with a 456 nm LED (50W max) for 3 minutes. Cell images were taken 48 hours after the LED irradiation event. As shown in Figure S7, cells only express the mCherry protein, not the GFP protein without light-irradiation. This is because the ‘clamped’ sgRNA-12 has no gene editing activity and the expression of GFP was completely quenched by the stop codon and the frame shift. However, 456 nm photo-irradiation cleaved the DEACM linker, activating the sgRNA-12. The Cas9 enzyme was therefore able to bind and cleave the designed DNA target sequence between the mCherry open reading frame and the GFP open reading frame. The cellular NHEJ pathway then generated random insertion or deletion which may destroy the stop-codon and correct the frame shift, resulting in the activation of the GFP expression. Notably, the ‘clamping’ of the sgRNA completely deactivated its gene editing activity. Therefore, we observed no leakiness of GFP expression in the non-irradiated cells.

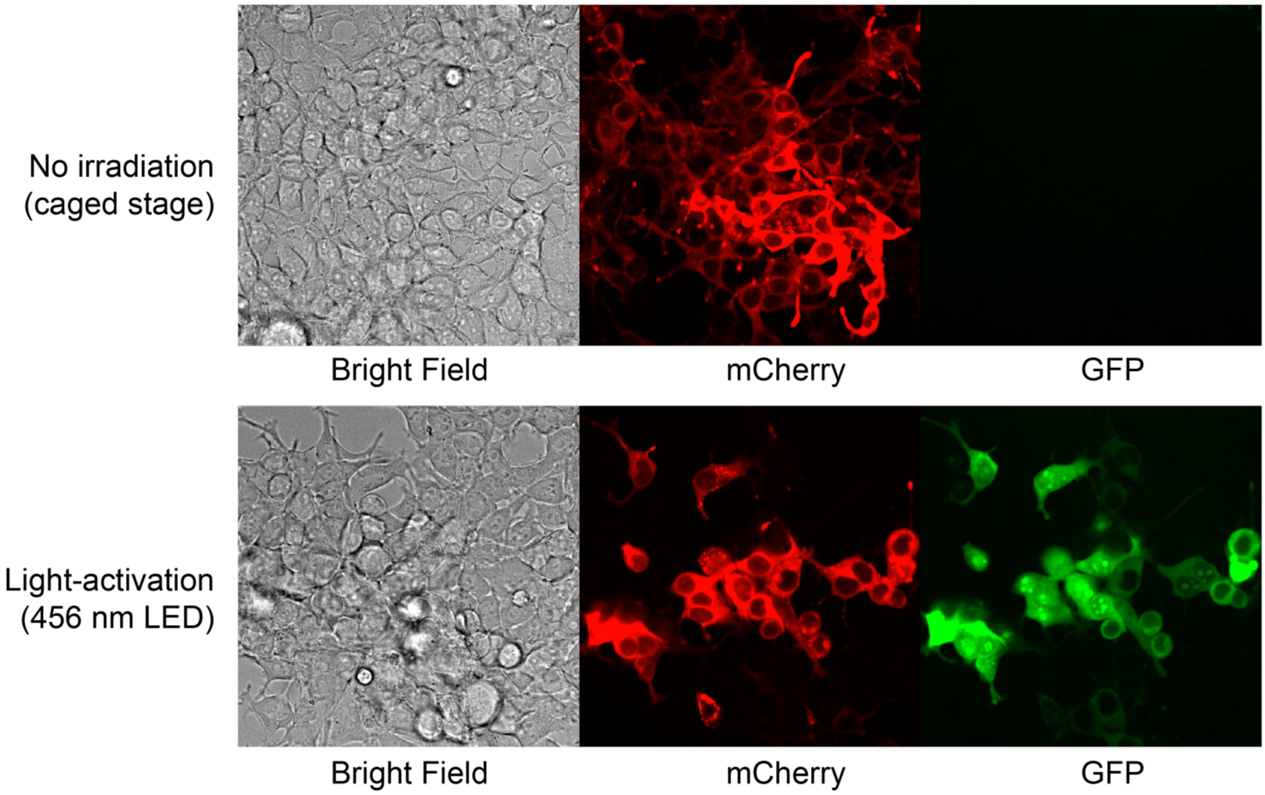

Figure S7. 456 nm LED photo-activation of gene editing using the surrogate EGFP reporter cell line. Without gene editing at the locus between the mCherry and GFP open reading frames, cells only expressed mCherry protein, not the GFP protein. Once Cas9-mediated gene editing happened at the designed locus, followed by the cellular NHEJ DNA repair pathway, the expression of the GFP protein was activated.

### In vitro binding assay of the ‘clamped’ sgRNA

We investigated the binding of the unmodified or ‘clamped’ sgRNA with the Cas9 protein (NEB, catalog number M0646T) in vitro. First, we designed a double-stranded 60 bp DNA substrate which contains the sgRNA-9 target sequence with the required NGG PAM. We used sgRNA-11 as the negative control non-targeting sgRNA, since sgRNA-11 did not target the 60 bp DNA substrate.

To test the binding the sgRNA and the 60 bp DNA substrate in vitro, 200 nM of Cas9 enzyme and 200 nM of sgRNA were incubated in 1X NEB buffer 3.1 for 15 minutes at room temperature. Next, 20 nM of DNA substrate was added into the pre-formed Cas9-sgRNA RNP and incubated at 37°C for 90 seconds. After the incubation, the reaction mixture was immediately placed on ice with the addition of 50% reaction volume of 50% glycerol to stop the reaction (Note: it is important to quench the reaction as quickly as possible.). Finally, the reaction was analyzed on 5% native TBE-PAGE (Figure S8). In Figure S8, lane 1 to lane 3 are the loading control of the sgRNA. The unmodified sgRNA-9 and sgRNA-11 were shown as multiple bands in the native TBE gel due to their secondary/ternary structures. Interestingly, the ‘clamped’ sgRNA-9 was shown as a single major band on the native gel. We reasoned it was because the intramolecular clamping rigidified the structure of the sgRNA. The formation of the sgRNA-9 Cas9 RNP was detected in lane 4. The formation of the Cas9-sgRNA-DNA ternary complex was only detected in lane 5 (unmodified sgRNA-9) and lane 6 (‘clamped’ sgRNA-9), but not in lane 7 (non-targeting sgRNA-11), demonstrating that the ‘clamped’ sgRNA-9 was also able to bind to the Cas9 as well as the target DNA in vitro.

| | 1 | 2 | 3 | 4 | 5 | 6 | 7 |
|---|---|---|---|---|---|---|---|
| 60 bp DNA substrate (20 nM) | - | - | - | - | + | + | + |
| Cas9 enzyme (200 nM) | - | - | - | + | + | + | + |
| sgRNA-9 (200 nM) | + | - | - | + | + | - | - |
| “clamped” sgRNA-9 (200 nM) | - | + | - | - | - | + | - |
| sgRNA-12 (200 nM) | - | - | + | - | - | - | + |

5% native PAGE
135 V 45 min

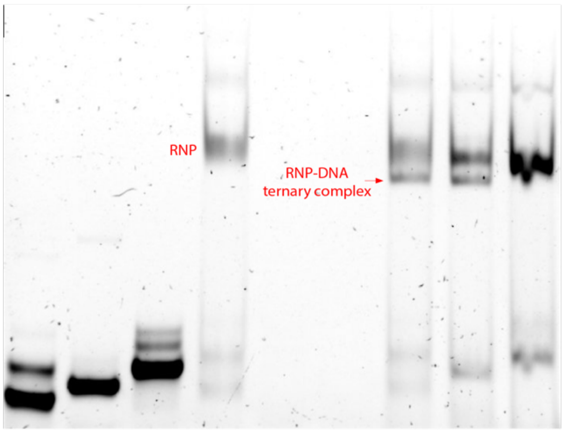

Figure S8. In vitro binding assay of unmodified or ‘clamped’ sgRNA and the DNA substrate.

### Spectrums

preQ1-PEG10-preQ1

H[1]NMR

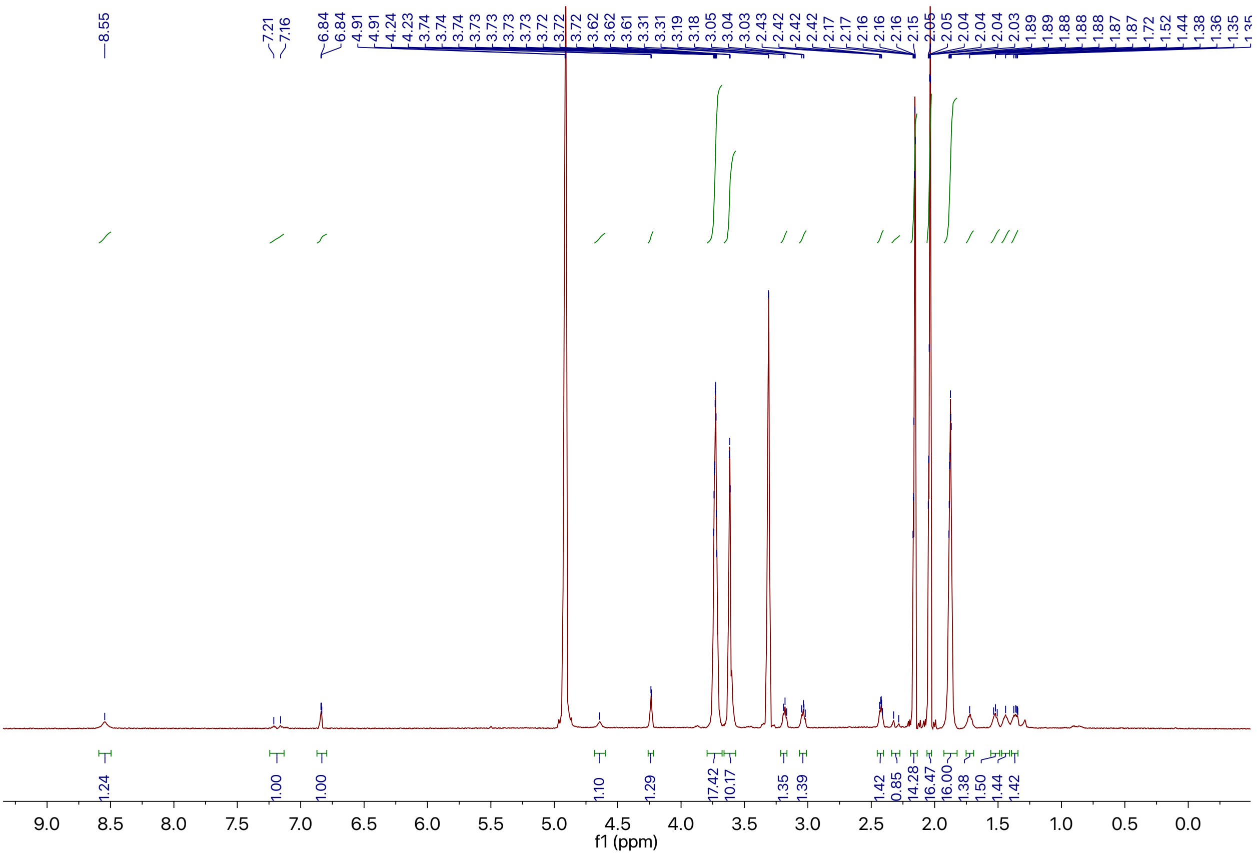

preQ1-PEG10-preQ1

C$^{13}$NMR

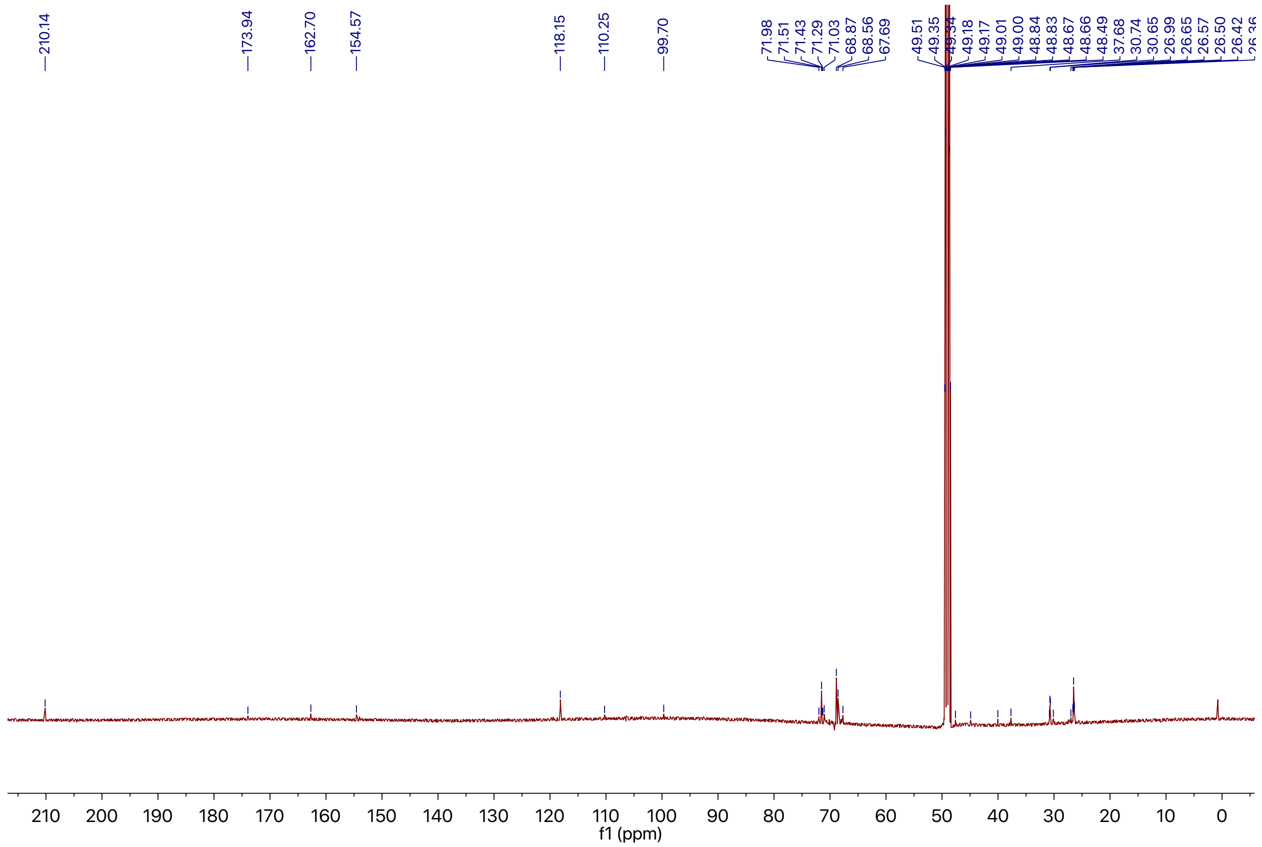

Alkyne-preQ1

H[1]NMR

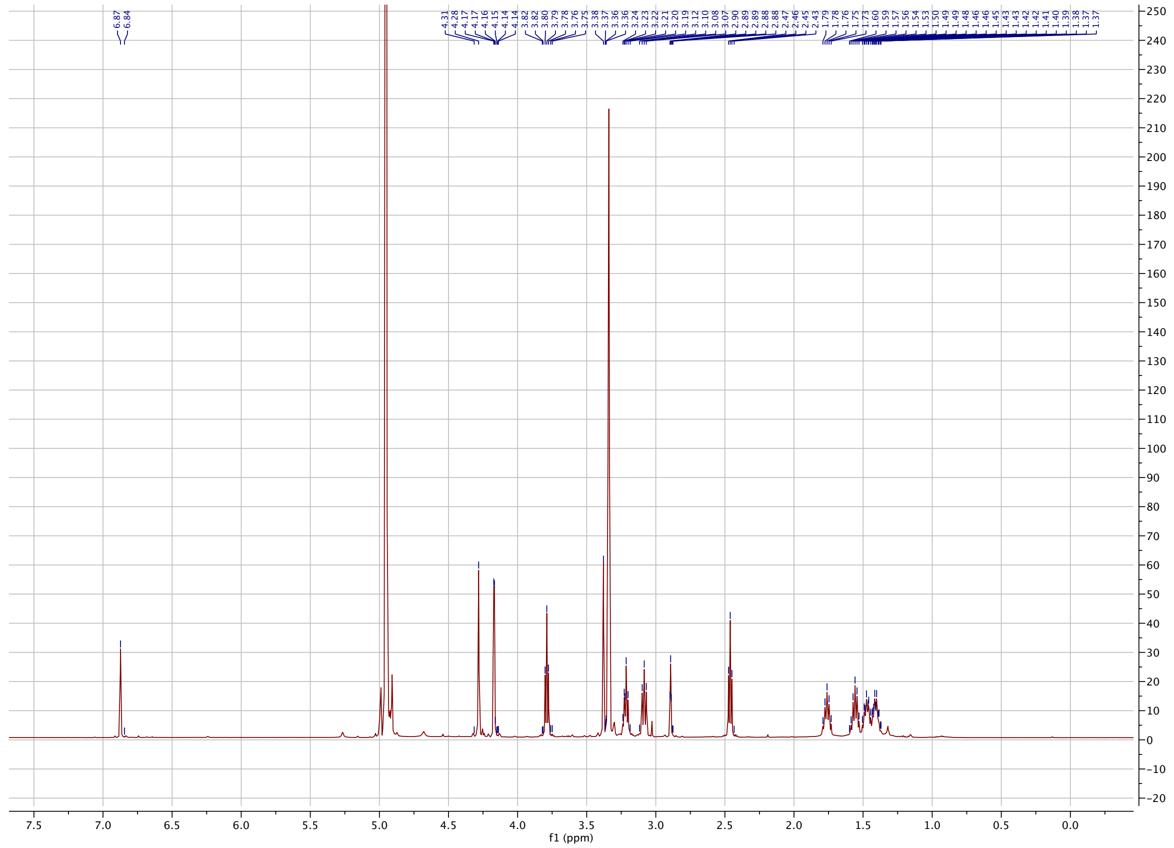

Alkyne-preQ1

$C^{13}$NMR

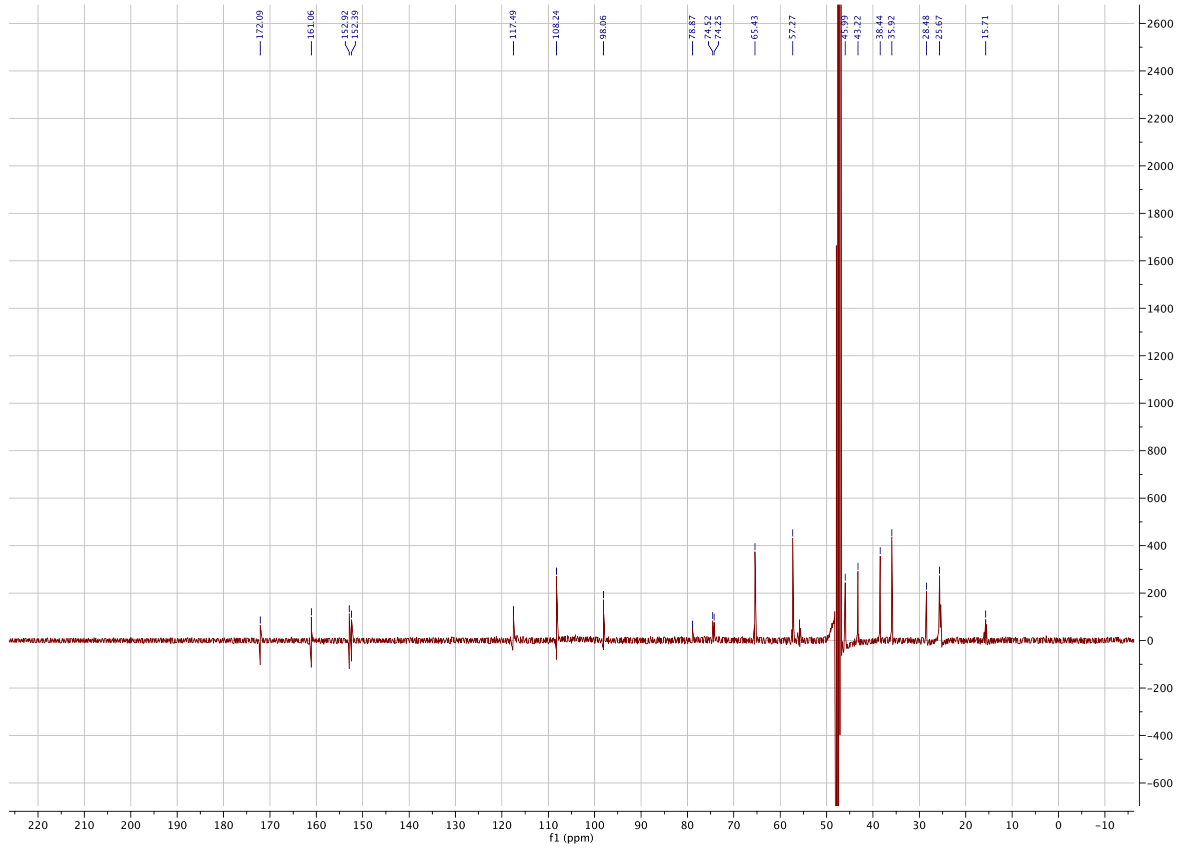

preQ1-PEG10-preQ1

HRMS

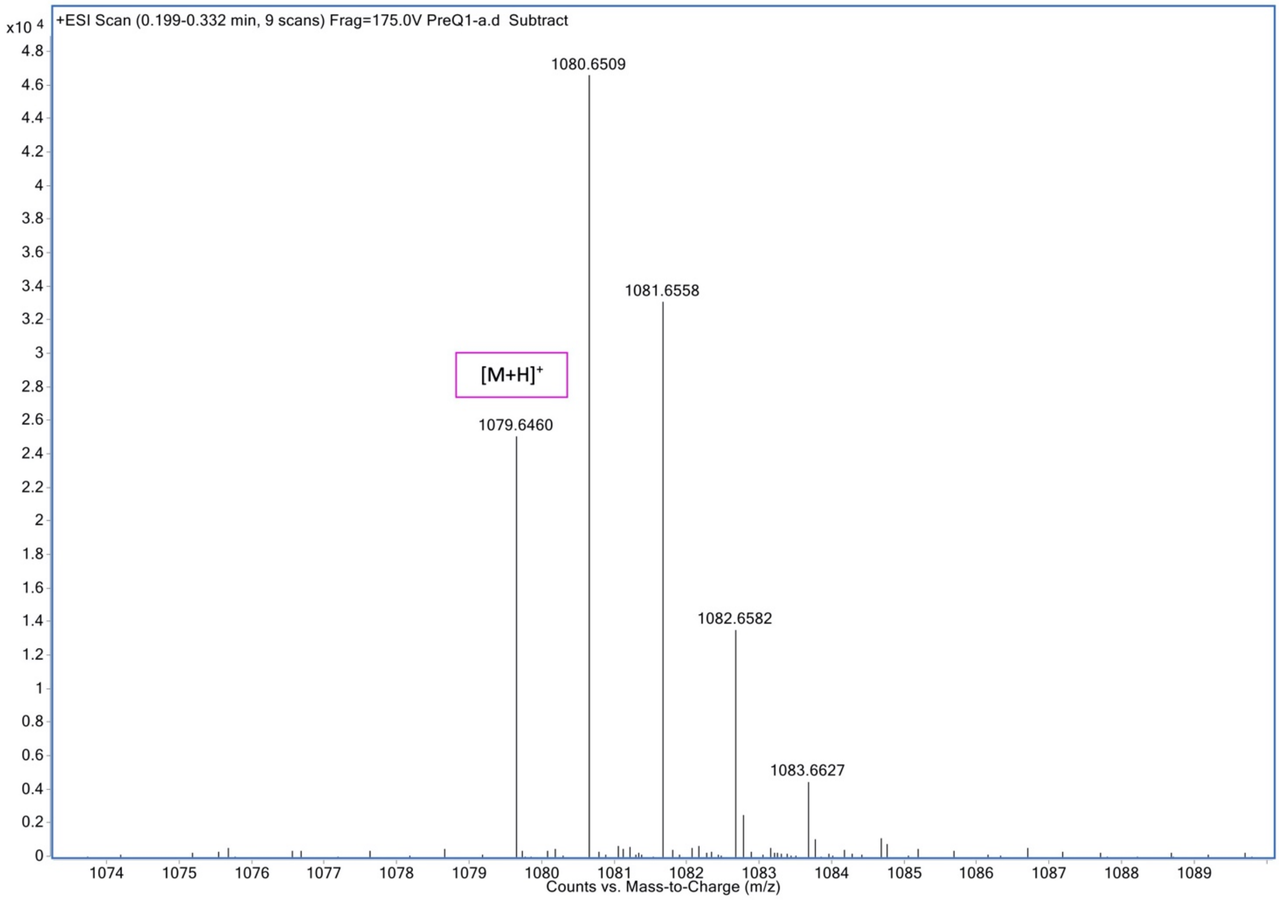

Alkyne-preQ1

HRMS

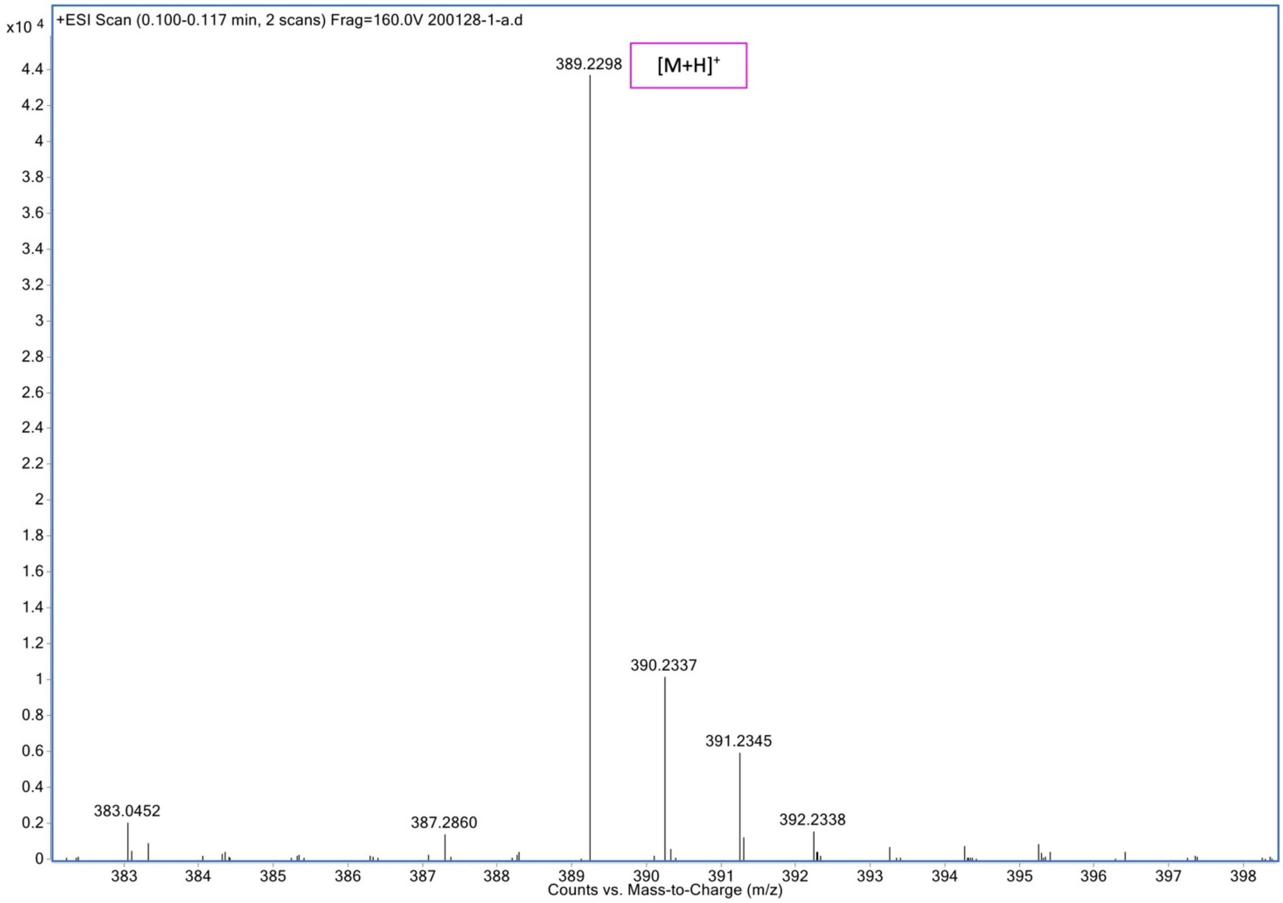

preQ1-DEACM-preQ1

HRMS

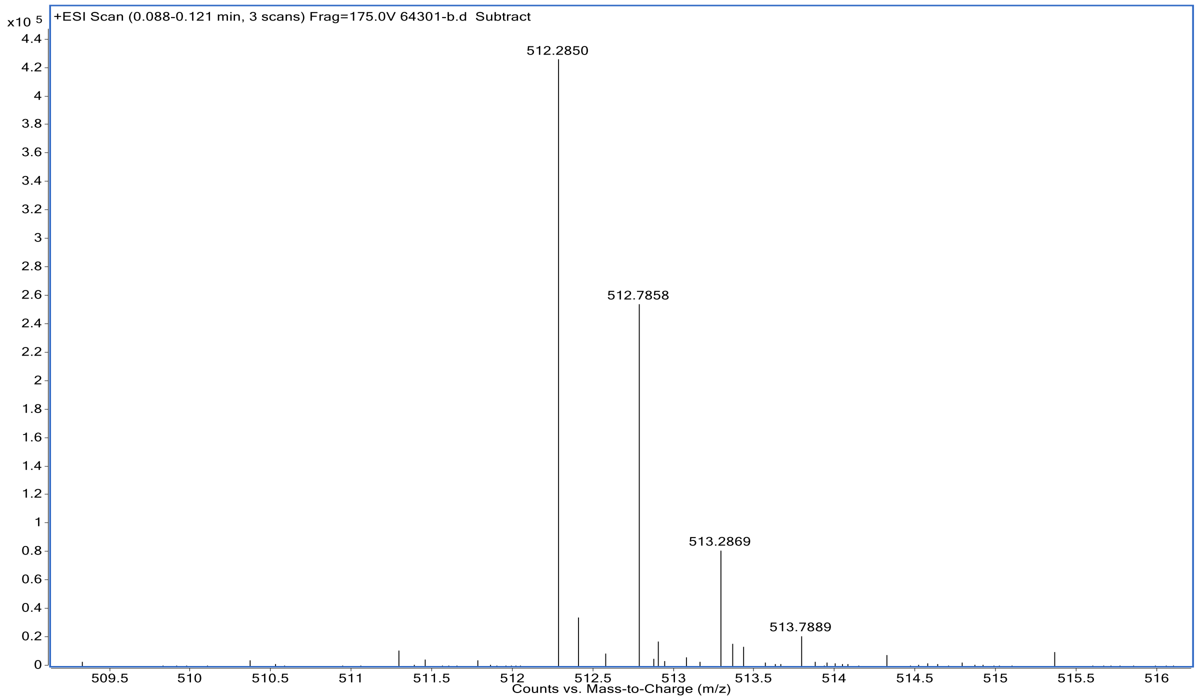

preQ1-NB-preQ1

HRMS

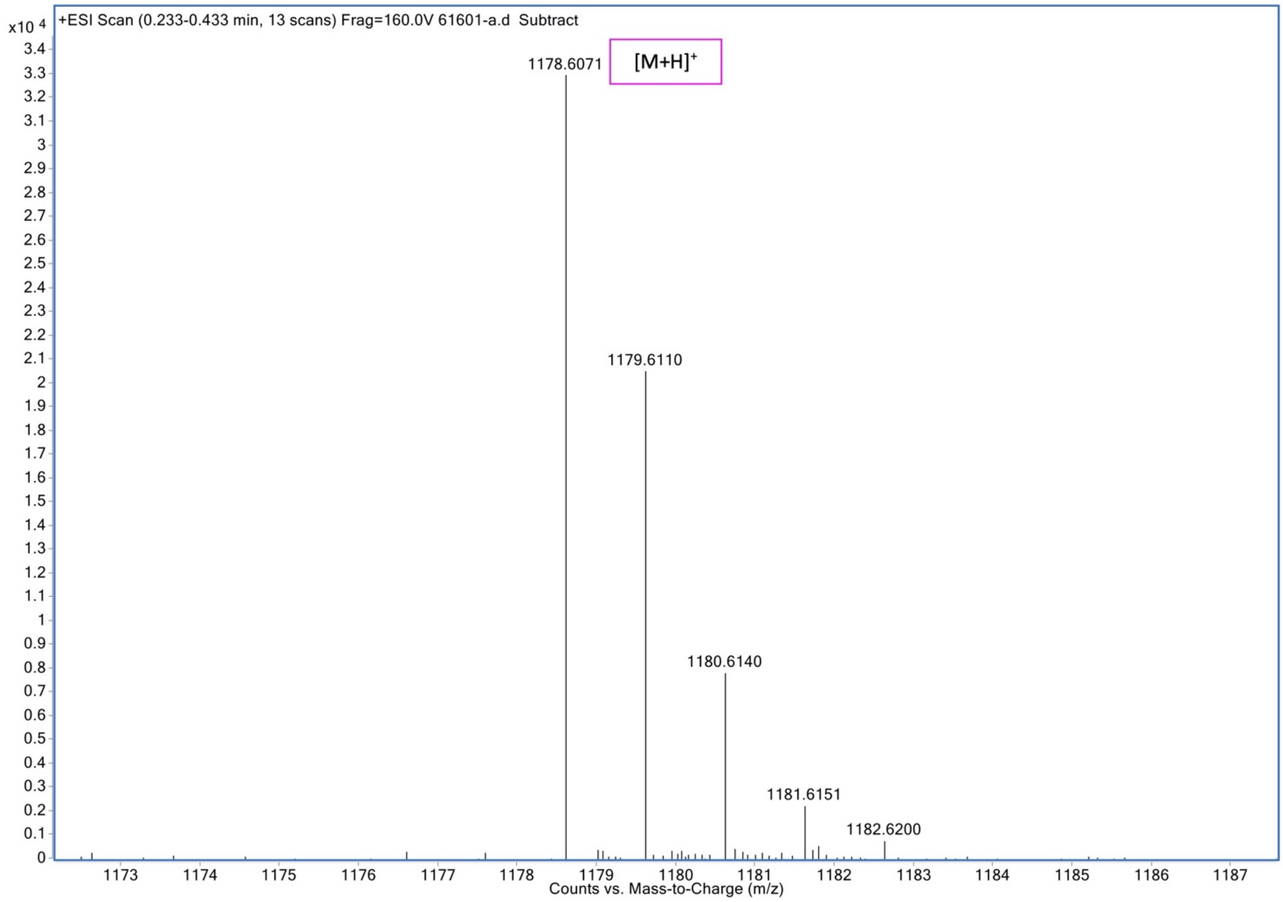
